# Ubiquitination modulates a protein energy landscape site-specifically with consequences for proteasomal degradation

**DOI:** 10.1101/843631

**Authors:** Emma C. Carroll, Eric R. Greene, Andreas Martin, Susan Marqusee

## Abstract

Cellular environments modulate protein energy landscapes to drive important biology, where small perturbations are consequential for biological signaling, allostery, and other vital processes. The energetic effects of ubiquitination are interesting due to its potential influence on degradation by the 26S proteasome, which requires intrinsically flexible or unstructured initiation regions that many known proteasome substrates lack. We generated proteins with natively attached, isopeptide-linked ubiquitin in structured domains to assess the energetic changes contributed by ubiquitin and how such changes manifest at the proteasome. Ubiquitination at sensitive sites destabilizes the native structure, and thereby increases the rate of degradation for substrates containing unstructured initiation regions. Importantly, this ubiquitination can even induce those requisite regions in well-folded proteins for proteasomal engagement. Our results indicate a biophysical role of site-specific ubiquitination as a potential regulatory mechanism for energy-dependent substrate degradation.

## Introduction

A protein’s function and folding is defined by its energy landscape, a term which encompasses all the accessible conformations, their relative populations, and the rates of interconversion. This energy landscape is determined by a protein’s amino-acid sequence and environment, and small changes in those parameters modulate this landscape. The phenotypic effects of these changes can range from undetectable to pathological^1–5^. Posttranslational modifications (PTMs) are one important environmental change that affect the energy landscape. Of the many PTMs that have been shown to affect protein structure and function^6^, the attachment of ubiquitin to lysine side chains is particularly interesting, as one of the most important roles for ubiquitination is to target proteins for degradation by the 26S proteasome.

The ubiquitin-proteasome system (UPS) is responsible for the majority of protein turnover in eukaryotic cells. Ubiquitin is an 8.5 kDa protein appended to other proteins (substrates) through an isopeptide bond between its C-terminus and the amino group of lysines in the substrate. Ubiquitin itself contains seven lysines, such that additional ubiquitin molecules can be added to form chains of different lengths, linkage types, and topologies. The 26S proteasome is the executor of the UPS, using ubiquitin receptors to selectively bind poly-ubiquitinated substrates and degrade them in an ATP-dependent manner. The degradation activity resides in the proteasome’s 20S core particle, whose proteolytic sites are sequestered inside a central cavity. Substrates are delivered to the 20S core particle largely through the 19S regulatory particle (RP), which caps one or both sides of the barrel-shaped 20S core. The RP recruits ubiquitinated substrates, mechanically unfolds them with its AAA+ (<u>A</u>TPase <u>A</u>ssociated with diverse cellular <u>A</u>ctivities) ATPase motor, and translocates the unstructured polypeptides into the core particle for cleavage into small peptides.

Although ubiquitination is best known for its association with proteasomal degradation, it is involved in a wide array of other cellular processes, including DNA repair, protein trafficking, endocytosis, and epigenetic regulation^7^. Therefore, the proteasome must carefully differentiate between ubiquitinated proteins that should be degraded and those ubiquitinated for other purposes. Failure of the proteasome to properly regulate substrate selection results in aberrant degradation, wasted energy, and collapse of proteostasis.

Diverse ubiquitin chain lengths and topologies, including mono-ubiquitination, induce proteasomal degradation both *in vitro* and *in vivo*^8–12^. For example, although the classic degradation signal consists of K48-linked poly-ubiquitin chains, K63-linked chains have a similar affinity for proteasomal ubiquitin receptors^10^ and mediate robust substrate degradation *in vitro*^13–16^. Thus, ubiquitin chain length, linkage type, and topology alone cannot fully explain proteasome substrate selection.

Conformational properties also play a role in determining whether a protein is degraded by the proteasome. In order to engage with the proteasomal AAA+ motor for unfolding and translocation into the core particle, a substrate needs to contain an unstructured initiation region long enough to enter the central pore and interact with conserved pore loops of the ATPase hexamer^17–19^. Nonetheless, a significant percentage of proteasome clients lack an obvious unstructured region in their native structure^20^. For some substrates, another AAA+ translocase, Cdc48 (also known as p97 or VCP), has been implicated in preparing them through partial or complete unfolding for subsequent proteasomal engagement^21–23^.

An exciting possibility is that ubiquitination itself can modulate the landscape and expose an unstructured region for degradation initiation by the proteasome. Molecular dynamics studies suggest that ubiquitination may destabilize proteins, principally through a decrease in substrate conformational entropy^24, 25^. If true, does this destabilization populate a proteasome-engageable unstructured region? However, purification of ubiquitin-conjugated substrates with native isopeptide bonds has been a challenging hurdle^26^, and the experimental characterization of ubiquitin-mediated changes in protein energetics has therefore been limited to artificial, non-physiological ubiquitin-attachment strategies. The potential energetic effects of substrate ubiquitination on proteasomal degradation thus remain completely unknown^27^.

Here, we developed a generalizable protocol to generate milligram quantities of homogeneously mono-ubiquitinated proteins. In this system, ubiquitin is attached via a native isopeptide linkage to a lysine within the folded region of a protein domain. Using several single-lysine variants of the small protein barstar, we show that mono-ubiquitination leads to local and global energetic changes that are site-specific and can lead to significant protein destabilization. Furthermore, these energetic modulations can affect proteasomal processing. Substrate variants destabilized by attached mono-ubiquitin display enhanced proteasomal degradation rates in the presence of an unstructured region for initiation. Importantly, in the absence of any intrinsically unstructured regions, ubiquitin-induced energetic changes can transiently expose flexible initiation regions, presumably by allowing access to high-energy, partially unstructured states that are engageable by the proteasome. Our data establish a connection between ubiquitin-induced changes in substrate energetics and proteasomal processing. We propose that modulation of substrate energy landscapes by site-specific ubiquitination can play a consequential role for substrate engagement and degradation by the proteasome.

## Results

### Generalizable Strategy for Site-Specific Ubiquitination on Substrate Lysines

Traditional spectroscopic studies of protein energetics and dynamics require large amounts (multiple milligrams) of homogeneous sample, yet such quantities are not feasible using established strategies of ubiquitin attachment^20, 26^. Furthermore, many approaches employed for artificial ubiquitin attachment require harsh chemical conjugation conditions and result in non-physiological linkages. Here, we used a biochemically reconstituted enzymatic ligation and deubiquitination strategy to overcome these technical obstacles and produce ubiquitin-substrate conjugates with native isopeptide bonds.

Substrate proteins were expressed as C-terminal fusions to maltose binding protein (MBP) with a connecting linker containing a PPPY recognition sequence for the yeast HECT E3-ubiquitin ligase, Rsp5^28^. Since substrates lacking MBP were less efficiently conjugated with ubiquitin, we believe MBP acts as a scaffold, helping to promote productive E3-substrate interaction, as previously described^29, 30^. In addition, the substrate domain contains a single cysteine for fluorescein-maleimide labeling (Fig. 1a).

**Fig. 1.**
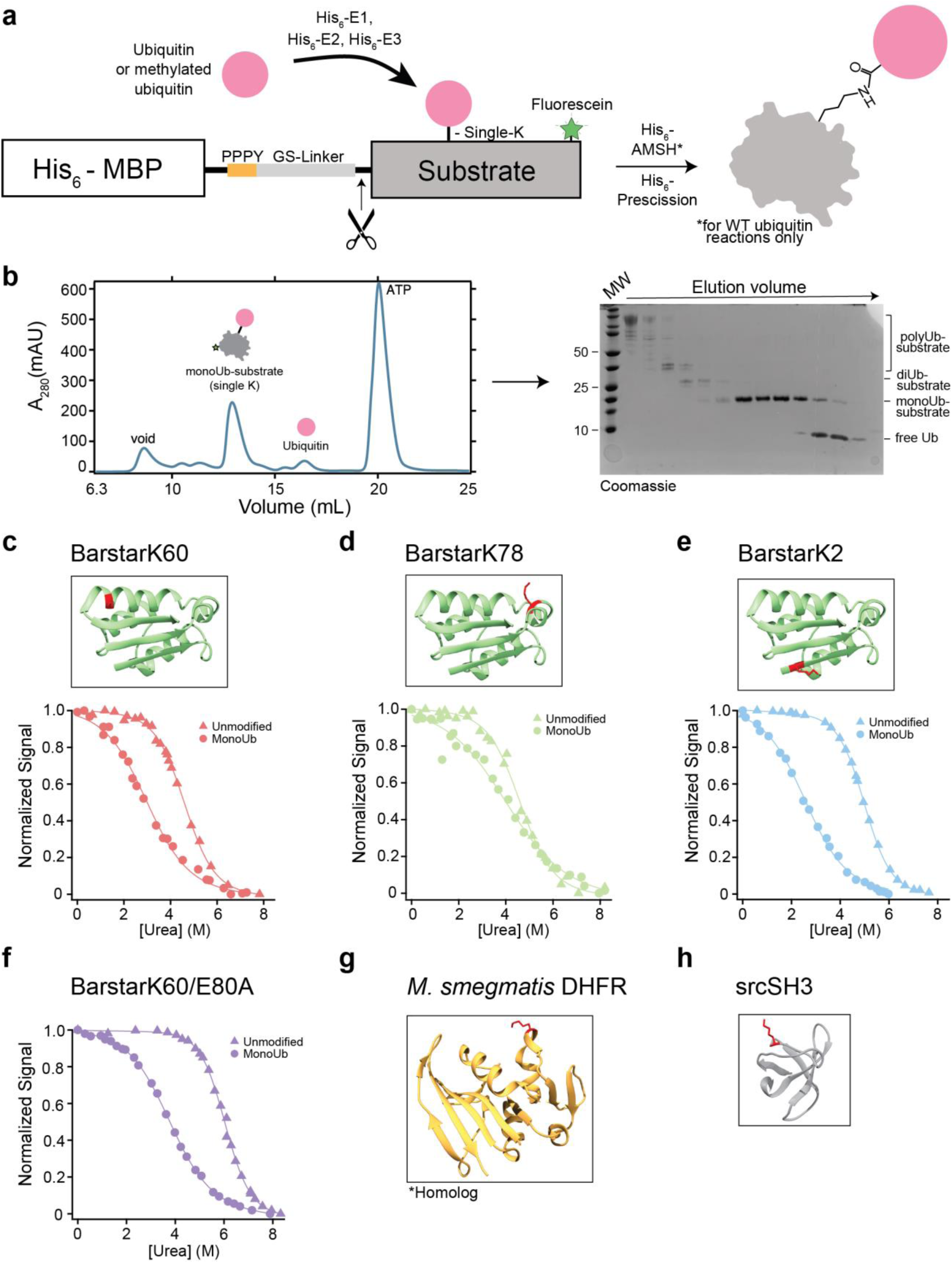
Generation of substrates with isopeptide-linked ubiquitin in structured regions and equilibrium unfolding studies. (**a**) Schematic of ubiquitination machinery and substrate design. A single-lysine substrate is fused to an N-terminal His-MBP scaffold with PPPY Rsp5 binding motif for enzymatic ubiquitination and subsequent purification. (**b**) Representative size exclusion trace of purified methylated monoUb-barstar and Coomassie-stained gel of selected size exclusion fractions. (**c-f**) Ribbon diagrams showing position of ubiquitinated barstar lysines in red (green: PDB: 1BTA) and urea melt of unmodified (triangles) and methylated monoUb-barstar (circles). (**g**) Ribbon diagram of *M. smegmatis* DHFR homolog from *M. tuberculosis* (orange: PDB: 1DG8) and *src*SH3 (grey: PDB: 1SRL) showing ubiquitinated lysine positions in red.

After purification, efficient *in vitro* poly-ubiquitination was achieved using a reconstituted system with mouse Uba1 as the ubiquitin-activating enzyme (E1), yeast Ubc4 as the ubiquitin conjugase (E2), and the yeast Rsp5 ubiquitin ligase (E3) (Fig. 1a). Treatment with the K63-specific deubiquitinase AMSH (<u>A</u>ssociated <u>M</u>olecule with the <u>SH</u>3 domain of STAM)^31^ collapses the heterogeneously poly-ubiquitinated substrates into lower molecular weight conjugates (Fig. 1a and Supplementary Fig. 1a-f). AMSH efficiently trims the Rsp5-generated, K63-linked ubiquitin chains, but is much slower in removing the proximal, substrate-attached ubiquitin moiety. We therefore varied the amount of AMSH and the length of the AMSH reaction before quenching with EDTA to accumulate the mono-ubiquitinated species. In addition, we generated mono-ubiquitinated substrates using methylated ubiquitin, which prevents chain formation and results in higher yield of modified protein for experiments requiring larger quantities. Although ubiquitin methylation has been observed to have various effects on the behavior and recognition of ubiquitin, we observed no difference compared to wild-type mono-ubiquitinated substrates in our biophysical measurements. For both approaches, a two-step subtractive Ni^2+^-NTA purification followed by size-exclusion chromatography was sufficient to purify the mono-ubiquitinated substrate to homogeneity (Fig. 1b). Using this generalizable method, we were able to attach ubiquitin to various single-lysine substrates (Fig. 1c-1h and Supplementary Fig. 1a-1f) and scale up the reaction to produce large enough quantities for spectroscopic measurements.

### Global Stability Assessment of Mono-ubiquitinated Substrates Demonstrates Site-Specific Effects

The ability to purify milligram quantities of homogenously mono-ubiquitinated proteins enabled us to determine global stability changes using traditional chemically-induced equilibrium unfolding monitored by intrinsic fluorescence. The observed fluorescence signal arises exclusively from tryptophan residues in our substrates, as wild-type ubiquitin is tryptophan-free. For these studies, we used a well-established model protein, barstar from *Bacillus amyloliquefaciens* (herein referred to as barstar), in which all except for one lysine were replaced by arginine to generate different single-lysine variants for site-specific ubiquitination. The position of the remaining lysine residue is denoted after the protein name (as in “barstarK60”, for a single-lysine barstar variant with the lysine at position 60).

Four single-lysine barstar variants were characterized: barstarK2, barstarK60, barstarK78, and barstarK60/E80A. We determined their equilibrium global stabilities in both unmodified and purified mono-methylubiquitinated forms by urea-induced chemical denaturation and fit the data using a two-state assumption and linear extrapolation (see Methods). The non-ubiquitinated versions of all four single-lysine variants display only minor destabilization compared to wild-type barstar^32^ (Fig. 1c-1f). In contrast, we observed dramatically different stabilities upon modification with mono-methylated ubiquitin, indicating site-specific effects (Fig. 1c-1f) (Table 1).

**Table 1.** Summary of thermodynamic values determined for all barstar variants and srcSH3 in this study. All reported errors represent S.E.M derived from curve fitting and propagated through all calculations. **^♯^**indicates that ΔG_unfolding_ values were calculated using a two state model with linear extrapolation. * indicates that ΔΔG_unfolding_ values were calculated from multiplying the C_m_ from the denaturation curves by the average *m*-value between the unmodified and mono-ubiquitinated proteins.

|  | BarstarK2 | BarstarK60 | BarstarK78 | BarstarK60/E80A | SH3 |
| --- | --- | --- | --- | --- | --- |
| $C_m$ unmodified<br>(M urea) | 4.97 +/-<br>0.28 | 4.41 +/-<br>0.55 | 4.65 +/-<br>1.19 | 6.02 +/-<br>0.38 | n.d. |
| $C_m$<br>monoUb<br>(M urea) | 2.52 +/-<br>0.16 | 2.51 +/-<br>0.34 | 4.24 +/-<br>1.05 | 3.72 +/-<br>0.26 | n.d. |
| $\Delta C_m$<br>(unmodified-<br>monoUb)<br>(M urea) | 2.45 +/-<br>0.32 | 1.90 +/-<br>0.65 | 0.42 +/-<br>1.59 | 2.30 +/-<br>0.46 | n.d. |
| $m$-value<sub>global</sub><br>unmodified<br>(kcal/mol M) | -1.06 +/-<br>0.04 | -0.96 +/-<br>0.08 | -0.89 +/-<br>0.17 | -1.16 +/-<br>0.05 | n.d. |
| $m$-value<sub>global</sub><br>monoUb<br>(kcal/mol M) | -0.59 +/-<br>0.02 | -0.68 +/-<br>0.07 | -0.48 +/-<br>0.09 | -0.70 +/-<br>0.04 | n.d. |
| $\Delta m$-value<sub>global</sub><br>unmodified-<br>monoUb<br>(kcal/mol M) | -0.47 +/-<br>0.05 | -0.28 +/-<br>0.11 | -0.41 +/-<br>0.19 | -0.47 +/-<br>0.06 | n.d. |
| $\Delta G_{unfolding}$<br>unmodified<sup>#</sup><br>(kcal/mol) | -5.27 +/-<br>0.20 | -4.25 +/-<br>0.38 | -4.16 +/-<br>0.74 | -6.99 +/-<br>0.31 | n.d. |
| $\Delta\Delta G_{unfolding}$<br>unmodified-<br>monoUb*<br>(kcal/mol) | -2.02 +/-<br>0.10 | -1.56 +/-<br>0.22 | -0.28 +/-<br>0.45 | -2.14 +/-<br>0.11 | n.d. |
| $\Delta G_{proteolysis}$<br>unmodified<br>(kcal/mol) | -2.72 +/-<br>0.02 | -3.24 +/-<br>0.11 | -2.72 +/-<br>0.06 | -3.52 +/-<br>0.01 | -2.70 +/-<br>0.12 |
| $\Delta G_{proteolysis}$<br>monoUb<br>(kcal/mol) | -1.66 +/-<br>0.10 | -2.12 +/-<br>0.06 | -2.66 +/-<br>0.09 | -2.56 +/-<br>0.08 | -2.38 +/-<br>0.23 |
| $\Delta\Delta G_{proteolysis}$<br>unmodified-<br>monoUb<br>(kcal/mol) | -1.07 +/-<br>0.11 | -1.12 +/-<br>0.13 | -0.06 +/-<br>0.15 | -0.96 +/-<br>0.12 | -0.32 +/-<br>0.10 |

Interestingly, all mono-ubiquitinated constructs show a small but notable decrease in *m*-value (the denaturant dependence of stability) compared to their unmodified counterparts (Table 1). *m*-values are known to correlate with the overall size of a protein or the non-polar surface area exposed during unfolding^33^, which may slightly change with the various ubiquitin attachments. Alternatively, these decreased *m*-values may indicate direct surface interactions with ubiquitin or a loss of two-state unfolding behavior, with the population of an unfolding intermediate^34, 35^. Because this questions the validity of the two-state assumption used to calculate ΔG_unfolding_, we report the midpoints of the denaturation curves (C_m_) for the unmodified and mono-ubiquitinated variants. Nevertheless, we used the average *m*-value of the unmodified/monoUb fits for each variant to determine an approximate ΔG_unfolding_ to provide a sense for the free energy changes associated with ubiquitination. BarstarK2 and barstarK60 were destabilized upon mono-ubiquitination, with C_m_ changes of 2.5 M and 1.9 M urea, respectively (Fig. 1c and 1e). A stabilized mutant of barstarK60, barstarK60/E80A, exhibited nearly identical net destabilization upon mono-ubiquitination, with a C_m_ decrease of 2.3 M urea (Fig. 1f). Conversely, barstarK78 displayed only marginal destabilization, with a C_m_ decrease of 0.42 M urea upon mono-ubiquitination (Fig. 1d). Using the average *m*-value of the unmodified/monoUb fits, these C_m_ changes correspond to ΔΔG_unfolding_ of −2.02 kcal/mol for barstarK2, −1.56 kcal/mol for barstarK60, −0.28 kcal/mol for barstarK78, and −2.14 kcal/mol for barstarK60/E80A (Table 1). Taken together, these results establish that the energetic effects of ubiquitin on a particular substrate can be highly site-specific, rather than being broadly destabilizing.

### Native State Proteolysis Reveals Energetics of Partial Unfolding

While the above results clearly demonstrate that mono-ubiquitin attached via a native isopeptide bond can alter a substrate’s global stability in a site-specific manner, the globally unfolded state is unlikely to be the most relevant fluctuation for proteasomal degradation. Under native conditions as in the cell, proteins sample partially folded conformations more frequently than the globally unfolded state. Furthermore, it is clear that the proteasome does not require global unfolding for successful substrate engagement.

To assess the population of partially unfolded states, we utilized a quantitative analysis of the susceptibility to a soluble protease, thermolysin^36, 37^. Because the active sites of soluble proteases typically require regions of ∼10-12 unstructured amino acids for productive proteolysis^36^, well-folded proteins under native conditions are proteolyzed via transient excursions to partially-folded, high-energy states (Fig. 2a). Typically, the lowest energy conformation that is competent for proteolytic cleavage (known as the “cleavable state”) will predominate. Because thermolysin has relatively poor affinity for its substrates (K_d_ ∼ 0.1-10 mM), proteolysis of the native state typically proceeds via an EX2-like kinetic regime, in which the proteolysis step itself, rather than the conformational change to the cleavable state, is rate-limiting^36^. As such, the observed rate of proteolysis is directly related to the free-energy change from the native state to this “cleavable” state (ΔG_proteolysis_) (Supplementary Fig. 2a). The ΔΔG_proteolysis_ for the same protein in two different states (i.e. unmodified and mono-ubiquitinated) can be reliably determined^36, 37^.

**Fig. 2.**
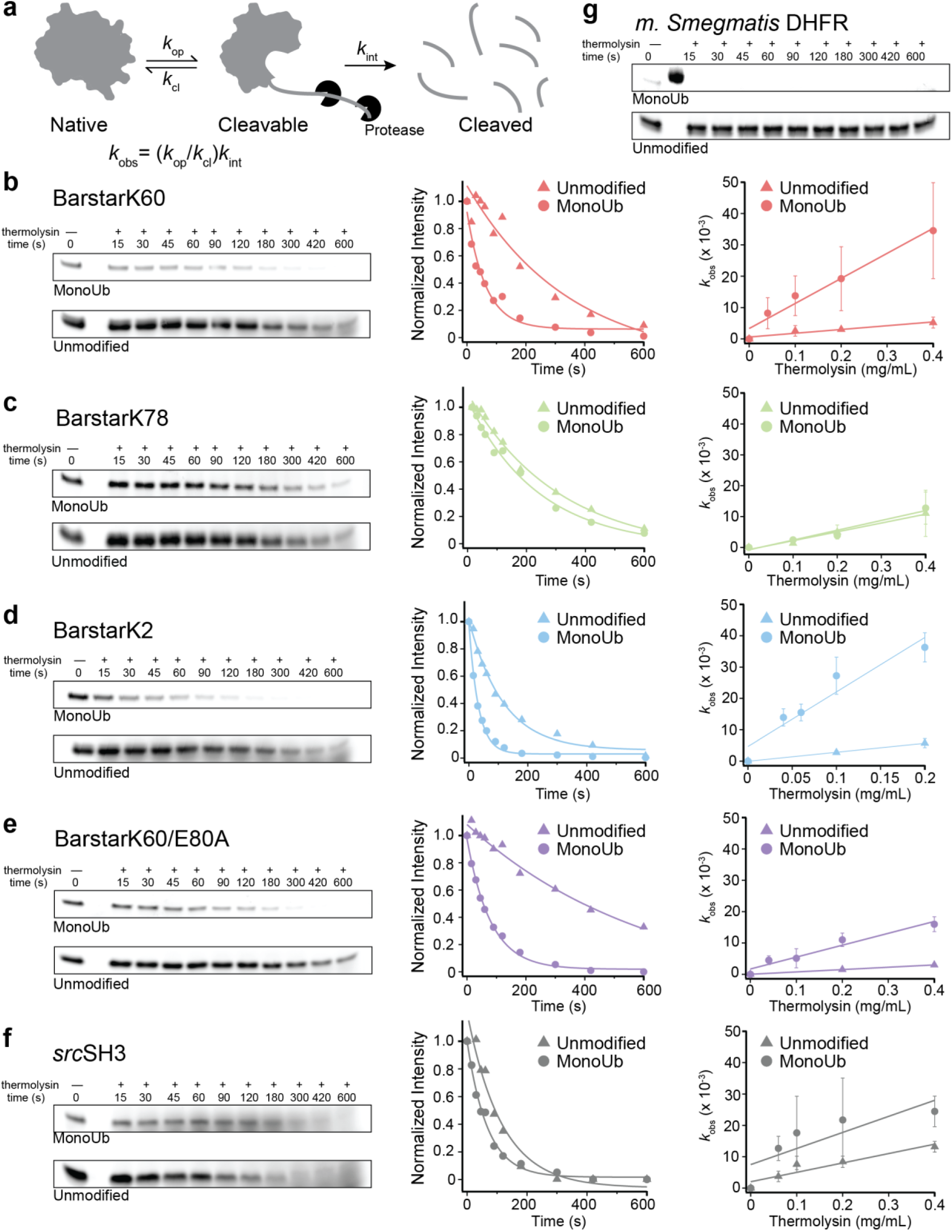
Native state proteolysis demonstrates the effects of mono-ubiquitination on the energetics of partial unfolding. (**a**) Under native conditions, well-folded proteins are proteolyzed via transient excursions to partially unfolded states. The observed rate of proteolysis, *k*_obs_, is proportional to the free energy of the conformational change from the native to partially unfolded state (ΔG_proteolysis_). (**b-f**) Representative native state proteolysis gels and quantified band intensities for indicated substrate proteins at 0.2 mg/mL thermolysin. ΔΔG_proteolysis_ upon mono-ubiquitination is calculated from the ratio of slopes of averaged *k*_obs_ (n≥ 3) against thermolysin concentration ranging from 0.04 to 0.4 mg/mL as indicated. Error bars represent the standard deviation of replicates. (**g**) Representative native state proteolysis gels for *M. smegmatis* DHFR at 0.2 mg/mL thermolysin.

We used this approach to measure the ΔG_proteolysis_ for unmodified and mono-ubiquitinated versions of all single-lysine barstar variants described above, as well as single-lysine *src*SH3 and *M. smegmatis* DHFR (wild-type is naturally single-lysine). AMSH concentration and reaction length were titrated to yield a mixture of both unmodified protein and protein modified with wild-type mono-ubiquitin, which allowed their direct comparison within the same proteolysis experiment.

Unmodified barstarK2, K60, and K78 exhibit nearly identical proteolysis kinetics (Supplementary Fig. 2b,c) and are clearly proteolyzed through sub-global unfolding fluctuations, as the calculated ΔG_proteolysis_ is less than ΔG_unfolding_ (Table 1). When comparing the ΔΔG_proteolysis_ values for the three pairs of unmodified and mono-ubiquitinated barstar variants, we observed a similar trend as for the global unfolding stabilities, with barstarK2 and barstarK60 showing significant changes in the population of the lowest energy cleavable state and a ΔΔG_proteolysis_ of −1.1 kcal/mol (Fig. 2b,d and Table 1). BarstarK60/E80A exhibited a similar net destabilization of −0.96 kcal/mol (Fig. 2e). Conversely, no destabilization was detected for barstarK78, indicating that there is no change in the free energy difference between the fully folded native state and the partially folded cleavable state upon mono-ubiquitination of this variant (Fig. 2c).

These variable effects on ΔG_proteolysis_ were recapitulated with other proteins. A single-lysine version of the *src*SH3 domain showed little destabilization with a ΔΔG_proteolysis_ of −0.32 kcal/mol (Fig. 2f). This is particularly interesting given that the *src*SH3 domain is smaller than the ubiquitin modification (64 aa vs 76 aa). In contrast, the naturally single-lysine *M. smegmatis* DHFR (159 aa) shows the most drastic destabilization upon mono-ubiquitination (Fig. 2g). The mono-ubiquitinated variant is completely proteolyzed within the dead time of the experiment (15 seconds), despite very little cleavage on this timescale for the unmodified DHFR species. Interestingly, monoUb-DHFR is still capable of binding methotrexate, albeit with greatly reduced affinity, suggesting that the native state is still populated (Supplementary Fig. 2d).

In sum, the changes in the energetics of the native state upon mono-ubiquitination are site-specific. The trends observed for the effect of mono-ubiquitin on global unfolding track with those of partial unfolding. These effects of ubiquitin on target-protein energetics can be accurately assessed using relatively small amounts of ubiquitin-conjugated sample.

### Ubiquitin-Mediated Modulation of the Energy Landscape Correlates with Proteasomal Degradation Rates

The ability of the proteasome’s AAA+ motor to unfold proteins is paramount to successful clearance of substrates and has been recently proposed as the rate-limiting step in the degradation process^16^. Therefore, we asked whether ubiquitin-mediated substrate destabilization conferred an increase to the proteasomal degradation rate. In order to directly compare mono-ubiquitinated substrates to their non-ubiquitinated counterparts, we used a system for ubiquitin-independent substrate delivery to the proteasome. In this system, a permutant of the bacterial SspB_2_ adaptor protein fused to the N-terminus of the Rpt2 ATPase in the proteasomal AAA+ motor recruits substrates that contain an ssrA sequence on a sufficiently long unstructured tail region for engagement^14^ (Fig. 3a and Supplementary Fig. 3a). All substrates delivered in this manner will be engaged equally, and thus, observed changes in overall degradation rate can be attributed to differences in substrate energetics. Proteasome-mediated degradation under single-turnover conditions (Supplementary Fig. 3b) was monitored by SDS-PAGE, and rates were determined based on both the disappearance of full-length substrate and the appearance peptide products (Fig. 3b and Supplementary Fig. 3c).

**Fig. 3.**
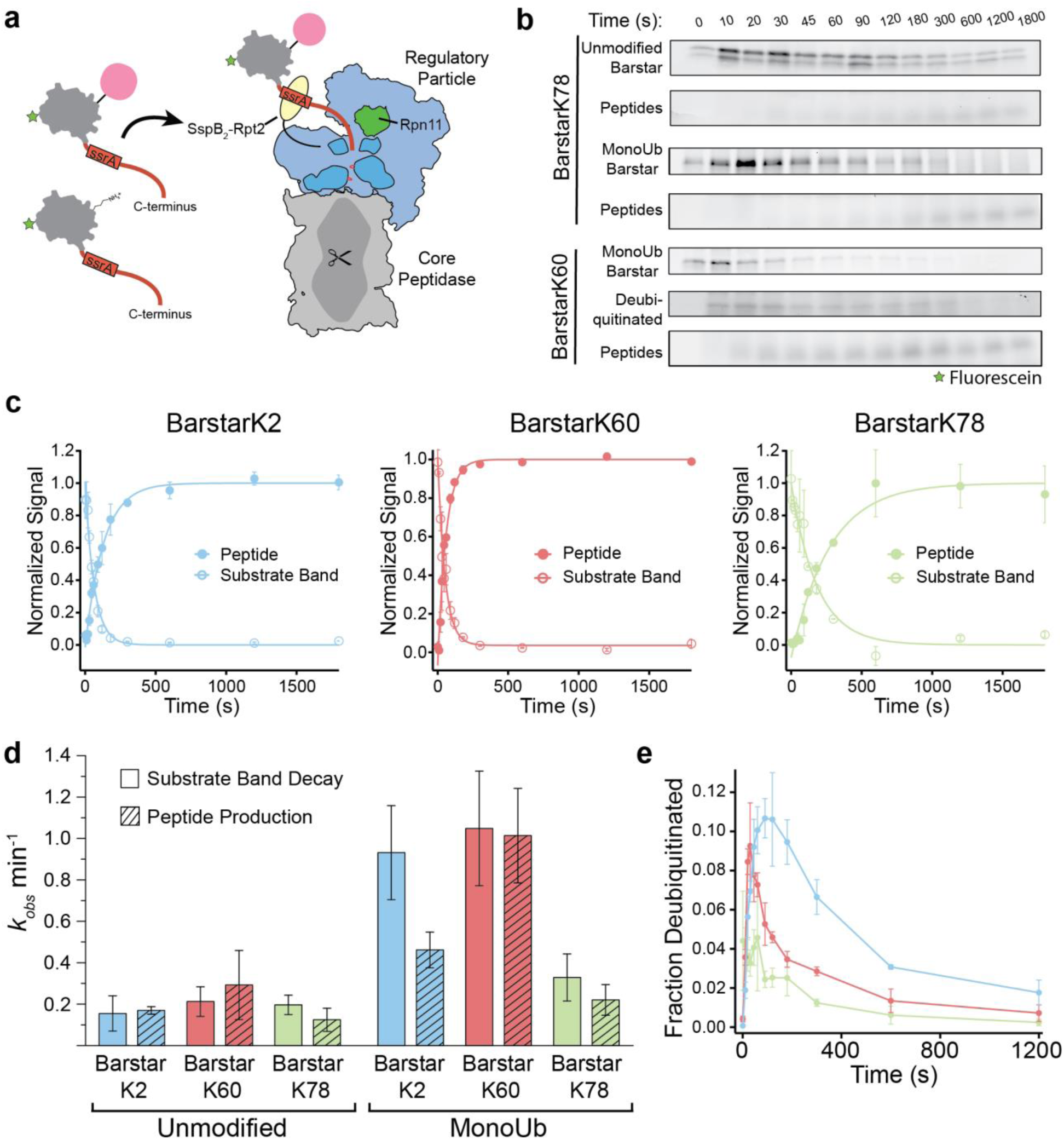
Mono-ubiquitin mediated substrate destabilization directly modulates degradation rate. (**a**) Schematic of ubiquitin-independent substrate delivery system where substrates are expressed with a flexible C-terminal tail containing an ssrA tag to direct delivery to the proteasome via an SspB_2_-dimer (yellow) fused to the base AAA+ ATPase. Core particle is represented in gray, regulatory particle in blue with Rpn11 in green, the AAA+ ATPase motor in dark blue, pore loops in red, substrate in gray with a green star representing fluorescein and red displaying the unstructured tail, and ubiquitin in pink. (**b**) Representative fluorescein-scanned SDS-PAGE gels showing disappearance of unmodified barstarK78 or mono-ubiquitinated (monoUb) barstarK78 and K60 with concomitant peptide production. The transient appearance of a deubiquitinated species for monoUb-barstarK60 is shown and quantified in **e**. (**c**) Normalized fractional signal plotted as averages with error bars representing standard deviation (n=3) of mono-ubiquitinated substrate band decay and peptide production. (**d**) Calculated rates and associated errors (S.E.M.) from **b** and **c**. (**e**) Fraction of total signal of deubiquitinated species plotted against time as averages and standard deviation (n=3) with line connecting points for visibility.

All ubiquitinated and non-ubiquitinated barstar variants were processed by the proteasome. As expected, all showed anti-correlated substrate depletion and peptide formation that were dependent on the presence of RP as well as ATP (Fig. 3b,c and Supplementary Fig. 3c,d). Degradation rates thereby correlated with the stability changes described above. Non-ubiquitinated barstar variants displayed similar degradation kinetics, with an observed rate (*k*_obs_) on the order of 0.1 - 0.3 min^-1^ (Fig. 3d). MonoUb-barstarK78 showed comparable kinetics, consistent with the negligible stability change upon ubiquitination of this variant (Fig. 3d). However, for the highly destabilized monoUb-barstar variants, degradation kinetics were substantially increased (*k*_obs_ = 1.04 min^-1^ for monoUb-barstarK60 and 0.93 min^-1^ for monoUb-barstarK2), suggesting that ubiquitin-mediated substrate destabilization increases the rate of unfolding by the proteasome.

For monoUb-barstarK60, we obtained similar results when following the substrate decay versus peptide production (Fig. 3c,d). For the monoUb-barstarK2 variant, however, these two processes were decoupled, with the mono-ubiquitinated species disappearing two times faster than the appearance of peptide products (Fig. 3c,d). This apparent decoupling may originate from differences in the temporal order of deubiquitination and unfolding. All variants showed a transient appearance of deubiquitinated species (Fig. 3e), accounting for ∼10% of the total substrate intensity for barstarK2 and barstarK60 at their peak. However, the deubiquitinated barstarK60 species was short lived (peaked at 30 s and negligible at 3 mins), while the barstarK2 species was observed to persist for ∼5 mins. Differences in the placement of ubiquitin relative to the substrate-engagement site (the C-terminal unstructured tail) may change the timing of ubiquitin cleavage relative to crossing the unfolding barrier. In the native barstar structure, the N- and C-termini are located in close proximity (Fig. 1c-e and Fig. 3a; PDB 1BTA). Engagement via the fused C-terminal tail may therefore place a ubiquitin attached to K2 in close proximity to the proteasome’s deubiquitinase (Rpn11), allowing deubiquitination immediately after engagement and before unfolding and translocation. If deubiquitination occurs prior to substrate unfolding, the destabilizing effect conferred by ubiquitin is lost, resulting in a slower rate of peptide production compared to the disappearance of the ubiquitinated substrate. Other ubiquitin sites (such as K60 or K78) might require substrate unfolding and translocation to occur first in order to position the ubiquitin-modified lysine for deubiquitination. These data therefore support the correlation between a substrate’s thermodynamic stability and its rate of proteasomal degradation, and extend this hypothesis to include ubiquitin attachment as a mode of site-specific destabilization of substrate proteins.

### Substrate Ubiquitination Can Induce a Proteasome-Engageable Unstructured Region

We next investigated the effect of ubiquitin-induced energetic changes on substrate engagement by the proteasomal AAA+ motor. Numerous biochemical studies have demonstrated the role of an unstructured initiation or engagement region^17–19^, yet a substantial fraction of cellular proteasomal substrates appear to lack such flexible segments^20^, begging the question of how their degradation is initiated. While other unfoldases, like Cdc48/p97 may generate disordered regions^21, 23, 38^, it is also possible that for some proteins ubiquitin-mediated conformational changes are sufficient to expose the obligate unstructured segments. To test this hypothesis, we poly-ubiquitinated our panel of single-lysine barstar variants (Ub_n_-barstar) and assayed the proteasome’s ability to recognize these substrates via its endogenous ubiquitin receptors and degrade them in an ATP-dependent manner (Fig. 4a). Native-state proteolysis experiments showed that these poly-ubiquitinated barstar variants have similar energetic profiles as the mono-ubiquitinated variants detailed above (Supplementary Fig. 4a).

**Fig. 4.**
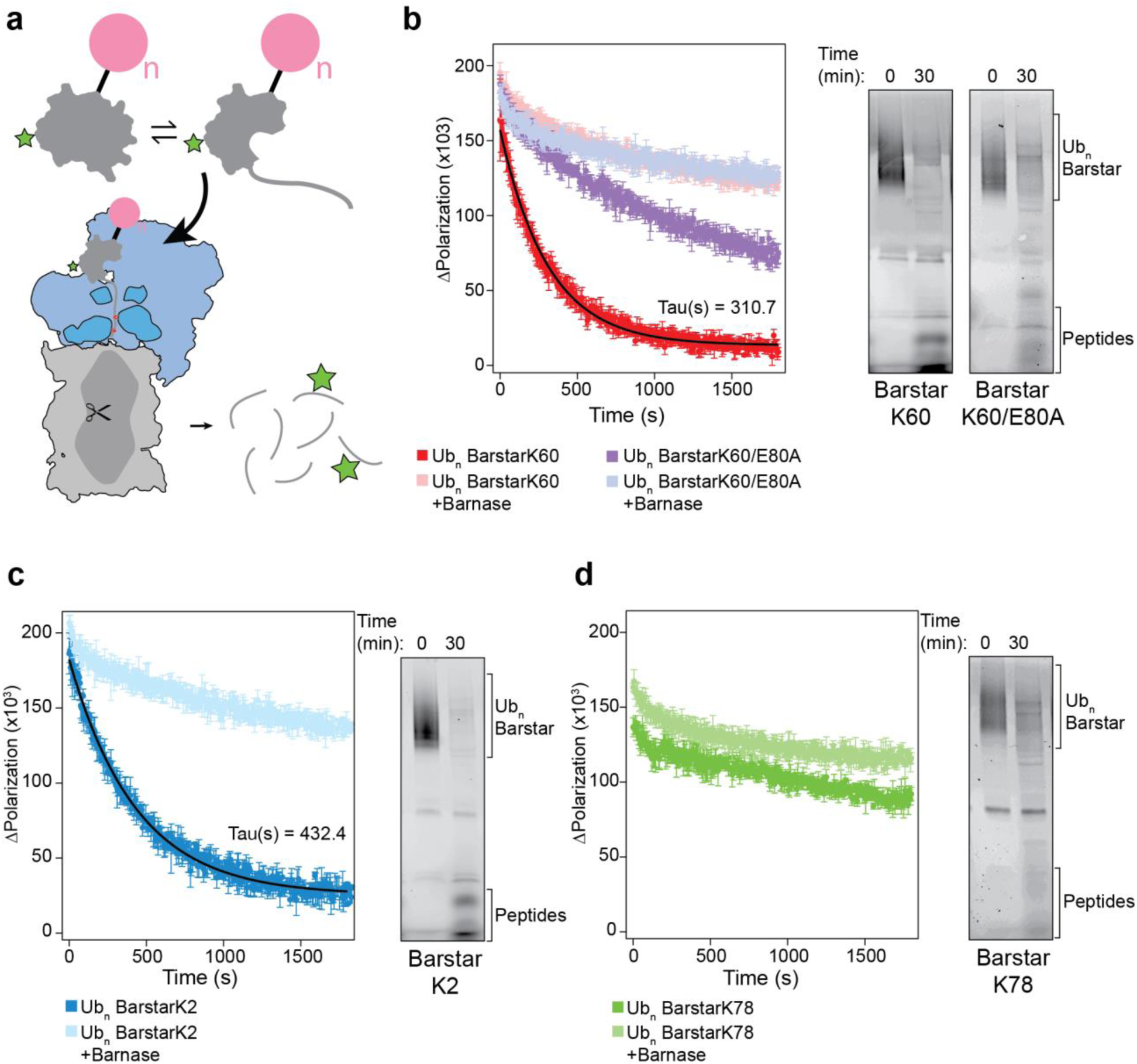
Ubiquitin destabilization of barstar is sufficient to release a proteasome-engageable unstructured region. (**a**) Schematic of degradation reaction showing Ub_n_-substrate lacking an unstructured region at equilibrium with a partially unfolded state, whereby the partially unfolded state is competent for proteasome engagement, translocation, and proteolysis. Core particle is represented in gray, regulatory particle in blue, the AAA+ ATPase motor in dark blue with pore loops in red, substrate in gray with a green star representing fluorescein, and ubiquitin in pink. Degradation can be monitored by fluorescence polarization as a change from large poly-ubiquitinated substrate to peptides. (**b-d**) Left: fluorescence polarization of Ub_n_-barstar single-turnover degradations in absence or presence of saturating barnase presented as averages and standard deviations (n=3) with line and calculated time constants (Tau). Right: fluorescein scan of SDS-PAGE gel from 30-minute endpoint of single-turnover Ub_n_-barstar degradations showing conversion of substrate to peptides and/or deubiquitinated species. Data also presented in Figure S4A.

Surprisingly, despite not harboring any obvious proteasome-engageable unstructured region, some poly-ubiquitinated single-lysine barstar variants were fully degraded into peptides by the 26S proteasome, whereas others were only slowly deubiquitinated (Supplementary Fig. 4b). Importantly, the kinetics of degradation into peptides depended on the site of the ubiquitination and correlate with the thermodynamics reported above. To gain a more quantitative understanding of the degradation kinetics, we labeled the barstar variants on Cys82 with fluorescein and monitored their cleavage into small peptides through changes in fluorescence polarization (Fig. 4a). Under single-turnover conditions (confirmed by varying the proteasome concentration, Supplementary Fig. 4c), Ub_n_-barstarK60 and Ub_n_-barstarK2 showed exponential degradation kinetics, with time constants of 310.7 s and 432.4 s, respectively (Fig. 4b,c). In contrast, Ub_n_-barstarK78 did not show any measurable degradation during these experiments, consistent with the hypothesis that site specific ubiquitin-mediated substrate destabilization determines whether the proteasome can engage an unstructured region (Fig. 4d). Furthermore, introducing the stabilizing mutation E80A to Ub_n_-barstarK60 substantially decreased degradation (Fig. 4b).

To further support our hypothesis that the ubiquitin-mediated modulation of barstar’s energy landscape is the principal determinant for its degradability by the proteasome, we added saturating concentrations of barnase, the high-affinity endogenous ligand of barstar, to these reactions (Supplementary Fig. 4d). In all cases, barnase ablated the degradation of substrates, while the remaining minimal decrease in fluorescence polarization could be attributed to minor degradation-independent deubiquitination (Fig. 4b-d). Addition of barnase has no effect on the degradation of a ubiquitinated titin substrate (FAM-Titin-I27^V13P,V15P^-35mer-tail substrate)^16^, confirming that the inhibition observed for the barstar variants was due to specific binding and stabilization of barstar folded state, rather than inhibitory interactions with the proteasome (Supplementary Fig. 4d).

In addition, we monitored degradation of the Ub_n_-barstar variants by the core particle alone to verify that robust degradation requires the entire 26S proteasome and includes ubiquitin recognition and ATP-driven unfolding/translocation. The core particle can only hydrolyze unstructured polypeptides that diffuse into its central chamber to access the proteolysis sites. Indeed, the core particle proteolyzes Ub_n_-barstarK2 and Ub_n_-barstarK60 only to a small extent and with low rates compared to the 26S holoenzyme (Supplementary Fig. 4c,e). Similar to the differences seen for the ATP-dependent degradation by the 26S proteasome, Ub_n_-barstarK78 did not show core-particle mediated degradation, and Ub_n_-barstarK60/E80A was cleaved by the core much more slowly than Ub_n_-barstarK60.

Unlike our observations with the ubiquitin-independent delivery system where we saw buildup of a deubiquitinated species for monoUb-barstarK2 (Fig. 3e), Ub_n_-barstarK2 did not populate a deubiquitinated species (Fig. 4c and Supplementary Fig. 4b). Because this substrate lacks the appended unstructured C-terminal tail, it must engage via a partially unfolded state, in which the ubiquitin attachment site may no longer be optimally positioned for Rpn11-mediated cleavage prior to substrate unfolding. Moreover, given that this variant is ubiquitinated very close to the N-terminus, it must be engaged by the proteasome C-terminal to the ubiquitination site. This is confirmed by our observation that inhibition of Rpn11 deubiquitination by *o*-phenanthroline did not inhibit degradation of this barstar variant, but inhibited all other variants (Supplementary Fig. 5a). For Ub_n_-barstarK2, the polypeptide between the ubiquitin-attachment point, K2, and the fluorescein-labeled Cys82 (80 residues) is long enough to span the distance between the entrance of the AAA+ pore and the proteolytic active sites (> 55 residues; Supplementary Fig. 5b). Rpn11-inhibited proteasomes can therefore move this substrate far enough into the 20S core for the fluorescein dye to be cleaved off, before translocation stalls on the K2-attached ubiquitin chain^39^.

### Ubiquitin-Induced Unfolding of an Engagement Region is Rate-Limiting for Degradation

The proteasomal degradation rates observed for poly-ubiquitinated barstar variants are notably lower than for barstar or other substrates with fused flexible tails^15, 16, 40^, suggesting that engagement of a spontaneously unfolding region represents the rate-limiting step for degradation. To probe this further, we turned to a proteasome variant, Rpn5-VTENKIF, whose mutations in the regulatory particle affect the conformational equilibrium of the proteasome and thereby hinder the insertion of flexible segments into the AAA+ pore, making engagement rate-determining even for moderately stable substrates with permanently unstructured tails^40^. Using this proteasome mutant, we see a three-fold (Ub_n_-barstarK2) and two-fold (Ub_n_-barstarK60) decrease in degradation rate (Supplementary Fig. 5c), suggesting that the slow degradation kinetics for our ubiquitinated, tail-less substrates are indeed determined by slow engagement and not unfolding by the proteasome. This leads to the interesting conclusion that for substrates without an easily accessible unstructured region, exposure of a flexible segment through spontaneous unfolding determines the rate of degradation, providing an alternative means of regulation for proteasomal targeting.

## Discussion

Clearance of damaged, misfolded, and regulatory intracellular proteins is paramount for sustaining life, and to a large extent catalyzed by the UPS. While substrate energetics have been known to critically affect the degradation of various substrates^16, 18, 41–43^, the influence of the substrate-attached ubiquitin itself has been elusive. Here, we provide evidence that ubiquitin can mediate substrate destabilization with direct consequences for proteasomal degradation. These effects can dictate both how fast a protein is degraded and whether a protein is susceptible to proteasomal degradation at all, thus providing an additional regulatory mechanism for clearance of a ubiquitinated substrate based on its conformational and energetic properties.

To carry out these studies, we developed a generalizable system to produce ubiquitin-modified single-lysine proteins with native isopeptide bonds (Fig. 1a). We were able to achieve efficient ubiquitination of several different single-lysine substrates, and we expect that this strategy will be useful to address a number of biological questions that are currently hampered by challenges in producing and purifying sufficient amounts of proteins carrying natively attached ubiquitin on structural domains^26^.

Our results clearly show that ubiquitin can affect a protein’s energy landscape with consequences for proteasomal degradation. There are no clear patterns regarding the region or type of secondary structure within the substrate that is energetically sensitive to the attachment of ubiquitin, nor are the effects correlated with the substrate size, as previously suggested^27^. It is reasonable to expect that the addition of a protein domain, such as ubiquitin, can alter the energetics and dynamics of a target protein in this manner. Biophysical studies of multidomain proteins have demonstrated that the stability of one domain can be modulated by the presence of another^44^. In differentially-linked polyubiquitin chains, the ubiquitin monomers themselves can have different thermodynamic and mechanical stabilities^45, 46^. Studies on N-terminal ubiquitin fusions and disulfide-linked ubiquitin attachments have reported small changes in the midpoints for thermally-induced unfolding depending on the modification^27^.

The changes we see in the energy landscape of our model substrates directly correlate with the measured changes in proteasomal degradation rates. By using tailed substrate variants and ubiquitin-independent delivery to the proteasome, we were able to selectively analyze the effects of ubiquitination on the mechanical unfolding by the AAA+ motor, and observed more rapid degradation for destabilized variants that reveal partially unfolded high-energy conformations. Perhaps more surprising is the fact that the energetic changes associated with poly-ubiquitination of small, cooperatively folded domains like barstar are sufficient to expose an unstructured region that meets the requirements for proteasomal engagement and thus enables substrate degradation. Hence, in addition to protein translocases like Cdc48/p97 that provide flexible regions for proteasomal engagement through partial or complete unfolding^21–23^, ubiquitination itself may play a role in destabilizing protein substrates and making them accessible for degradation by the proteasome.

Consistent with this concept, we found that stabilizing the protein via ligand binding (such as in barstar:barnase) inhibits proteasomal processing. The engagement of these substrates appears to be rate-limiting and modulated directly by the accessibility of partially-unfolded, proteasome-engageable states. Thus, the overall context of the ubiquitinated protein with respect to cellular environment, binding partners, and perhaps other stabilizing or destabilizing PTMs can influence whether a ubiquitinated substrate is actually degraded.

Based on our results, we can build a model for the effect of ubiquitin-mediated, site-specific changes in protein energy landscapes on proteasomal degradation (Fig. 5), in which: 1) a protein may or may not be engaged by the proteasome based on its altered energetics, and 2), the speed with which ubiquitinated substrates are degraded is related to the extent of ubiquitin-induced destabilization. Both aspects of proteasomal turnover are directly modulated by the increased sampling of partially-unfolded states and further influenced by other factors, such as stabilizing mutations or deubiqutination prior to substrate unfolding, either at the proteasome by Rpn11 or by a host of cellular deubiquitinases^47^.

**Fig. 5.**
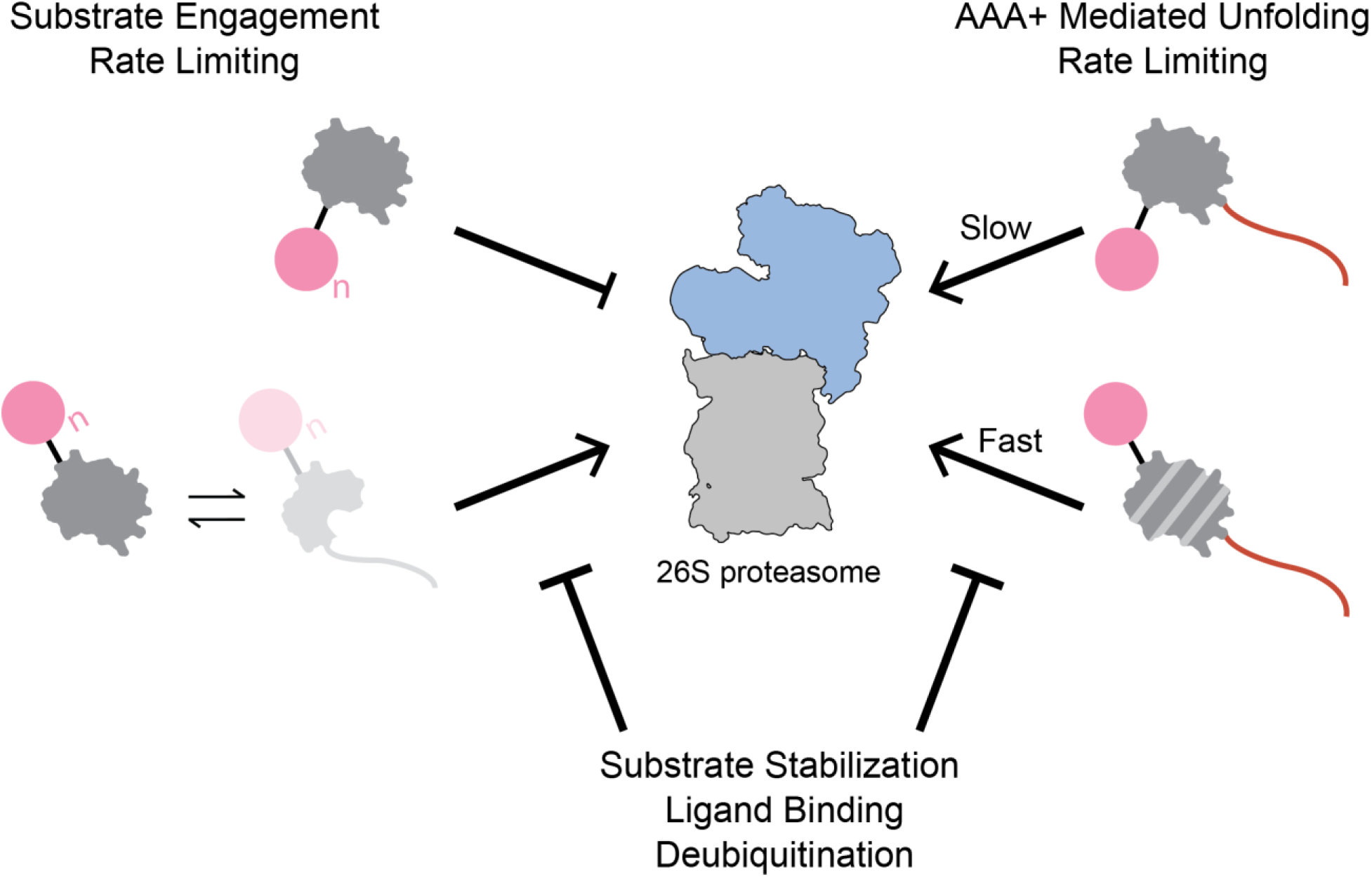
Model of the consequences of site-specific, ubiquitin-induced energy landscape modulation for proteasomal degradation. If ubiquitin-modification occurs on a non-sensitive structured lysine, as in barstarK78, the target protein does not populate a partially-unfolded, proteasome-engageable conformation to an extent that allows for successful degradation. If ubiquitin-modification occurs on a sensitive lysine, as in barstarK2 and barstarK60, the otherwise-tailless target proteins are sufficiently destabilized to populate partially-unfolded, proteasome-engageable conformations and are successfully degraded by the proteasome. The observed degradation kinetics thus appear dependent on the changes to the protein energy landscape upon ubiquitination. When substrates contain a proteasome-engageable tail, ubiquitination at sensitive lysine positions allows for substantially faster degradation kinetics, while degradation kinetics of substrates with non-destabilizing ubiquitinations remain essentially unchanged. Successful proteasome engagement and degradation of ubiquitin-destabilized substrate proteins can be substantially slowed or blocked completely by a number of energetically stabilizing events, including deubiquitination, addition of a binding partner/ligand, or stabilizing mutation.

This model has implications for a number of different processes, including the engineering of substrate degradation via Proteolysis Targeting Chimeras (PROTACs)^48^. PROTACs are synthetic molecules containing two moieties, a ligand binding the target protein to be degraded and another ligand with affinity for an E3 ubiquitin ligase that facilitates ubiquitination of the target. The linker length between the two ligands has been found to strongly affect whether the target protein is degraded^49^, likely because it determines which lysines of the target are ubiquitinated in a manner that facilitates delivery to the downstream processing enzymes, i.e. Cdc48/p97 and the proteasome^38, 50^ but also possibly depending on whether ubiquitination at these lysines destabilizes the target. Additionally, it was observed that non-specific ligands promiscuously binding to 50-100 protein kinases facilitate the degradation of only a small subset of these kinases^51, 52^, which also could be due to which lysines are ubiquitinated on the different targets and whether these ubiquitinations act sufficiently destabilizing to allow degradation.

While it is clear that ubiquitination has site-specific effects on the energy landscape, the mechanisms for ubiquitin-induced destabilization and the population of partially unfolded conformers remains unknown. Computational studies have postulated that the destabilization is conferred by ubiquitin via a decrease in the substrate’s overall conformational entropy^25^. In addition, NMR studies of the protein FKBP12 with chemically conjugated ubiquitin demonstrated increased backbone flexibility^27^. These findings could be rationalized by an entropic pulling model whereby a highly stable protein fold, like ubiquitin, attached to a substrate through a native isopeptide bond with many degrees of translational and rotational freedom can provide a net pulling force from the site of ligation^53, 54^.

Alternatively, the energetic modulation may arise from direct interactions between the ubiquitin and the substrate. Ubiquitin itself has multiple exposed hydrophobic patches, one near Ile44 and another at Ile36, which could potentially stabilize any exposed hydrophobic residues on a partially unfolded substrate. The Ile44 hydrophobic patch is known to interact with PCNA when in an N-terminal fusion^55, 56^ and be responsible for the inter-ubiquitin interactions that give K48-linked ubiquitin chains their compact conformation^57–59^. Ubiquitin also contains an acidic patch that has been found to electrostatically interact with some target proteins^60^. How exactly ubiquitin destabilizes the substrate protein remains unknown and will have to be further investigated.

Cellular proteostasis relies upon careful regulation of protein degradation via the UPS, and the consequences of aberrant degradation are severe. Here, we present evidence that ubiquitin directly modulates a protein’s conformational energy landscape, and these energetic changes play a pivotal role in regulating both substrate selection and the kinetics of degradation by the 26S proteasome. We conclude that ubiquitin signaling and proteasomal degradation overall are dependent on the biological and biophysical contexts of individual ubiquitinated proteins. A full understanding of the energetic effects contributed by a particular ubiquitination event is therefore crucial for building a complete model of how ubiquitin-mediated signals are transduced *in vivo*. We hope the tools developed herein can facilitate addressing these questions and, further, be used to expand our model of the biophysical factors governing ubiquitin-mediated signaling.

## Acknowledgements

We thank all members of the Marqusee and Martin labs for helpful discussions. We also thank Dr. Brendan Maguire and Dr. Ken Dong for assistance with protein purification and helpful troubleshooting expertise. S.M. acknowledges support from the US National Institutes of Health (grant R01 – GM050945) and the Chan Zuckerberg Biohub. A.M. is an investigator of the Howard Hughes Medical Foundation and acknowledges support from the US National Institutes of Health (grant R01-GM094497).

## Author Contributions

E.C.C. and E.R.G. performed the experiments and analyzed data. All authors contributed to experimental design, data interpretation, and manuscript preparation.

## Online Methods

### Protein Purification

#### Preparation of substrate proteins

*E. coli* BL21 Rosetta 2 (DE3) cells were transformed with either pEC072 (single-lysine *src*SH3), pEC074 (*M. smegmatis* DHFR), pEC076 (barstarK2), pEC062 (barstarK60), pEC081 (barstarK60/E80A), or pEC059 (barstarK78). Cells were then grown in 2 L LB Broth (Fisher) to 0.4 < OD_600_ < 0.8 and induced with 1 mM IPTG for 3 hours at 37°C. Bacteria were then pelleted and resuspended in 50 mM HEPES pH 7.0, 150 mM NaCl, 0.5 mM TCEP supplemented with 1X Halt^TM^ protease inhibitor cocktail (Thermo) and benzonase (Novagen). Resuspended cells were lysed by sonication and the lysate was clarified by centrifugation at 13,000 rcf, 4°C, 30 minutes. The substrate was first purified by Ni^2+^-NTA affinity chromatography using its N-terminal His_6_ tag. Clarified lysate was allowed to batch bind to HisPur^TM^ Ni^2+^-NTA resin (Thermo) washed with 50 mM HEPES pH 7.0, 150 mM NaCl, 25 mM imidazole, 0.5 mM TCEP and eluted with 50 mM HEPES pH 7.0, 150 mM NaCl, 500 mM imidazole, 0.5 mM TCEP. Concentration of protein in the eluate was then measured using UV-Vis absorption at 280 nm. Eluate was then labeled for 2 hours at room temperature with 5X molar excess fluorescein-maleimide dye (Thermo). The labeling reaction was quenched with 10X molar excess DTT and unreacted dye was removed using a S200 16/60 size exclusion column (GE) pre-equilibrated with 25mM HEPES pH 7.5, 150 mM KCl, and 15 mM MgOAc. Peak corresponding to the labeled, full length His-MBP substrate was collected, and quantified by UV-Vis absorption at 280 nm and 495 nm according to the manufacturer’s instructions before addition of 10% glycerol and flash freezing to store at −80°C for future use.

#### Preparation of substrate proteins with C-terminal ssrA tag/cyclin B engageable tail

*E. coli* BL21 Rosetta 2 (DE3) cells were transformed with either pEC098 (barstarK2), pEC093 (barstarK60), pEC097 (barstarK78). Cells were then grown in 2 L LB Broth (Fisher) to 0.4 < OD_600_ < 0.8 and induced with 1 mM IPTG for 3 hours at 37°C. Bacteria were then pelleted and resuspended in 50 mM HEPES pH 7.0, 150 mM NaCl, 0.5 mM TCEP supplemented with 1X Halt^TM^ protease inhibitor cocktail (Thermo) and benzonase (Novagen). Resuspended cells were lysed by sonication and the lysate was clarified by centrifugation at 13,000 rcf, 4°C, 30 minutes. The substrate was first purified by Ni^2+^-NTA affinity chromatography using its N-terminal His_6_. Clarified lysate was allowed to batch bind to HisPur^TM^ Ni^2+^-NTA resin (Thermo) washed with 50 mM HEPES pH 7.0, 150 mM NaCl, 25 mM imidazole, 0.5 mM TCEP and eluted with 50 mM HEPES pH 7.0, 150 mM NaCl, 500 mM imidazole, 0.5 mM TCEP + 1X Halt^TM^ protease inhibitor cocktail. Eluate was diluted 1:2 with 50 mM HEPES pH 7.0, 150 mM NaCl, and 0.5 mM TCEP and 5 mM EDTA was added. Eluate was then batch bound to *Strep*-Tactin Superflow Plus resin (Qiagen), washed with 50 mM HEPES pH 7.0, 150 mM NaCl and eluted with 50 mM HEPES pH 7.0, 150 mM NaCl, 2.5mM desthiobiotin (Sigma). Eluate was labeled with 5X molar excess fluorescein-maleimide dye (Thermo). The labeling reaction was quenched with 10X molar excess DTT and unreacted dye was removed using a S200 16/60 size exclusion column (GE) pre-equilibrated with 25mM HEPES pH 7.5, 150 mM KCl, and 15 mM MgOAc. The peak corresponding to the labeled, full length, labeled His-MBP substrate was collected, and quantified by UV-Vis absorption at 280 nm and 495 nm according to the manufacturer’s instructions before addition of 10% glycerol and flash freezing to store at −80°C for future use.

#### Preparation of Ubiquitin

*E. coli* BL21 Rosetta 2 (DE3) cells were transformed with pEC086. Cells were then grown in 2 L LB Broth (Fisher) to 0.4 < OD_600_ < 0.8 and induced with 1 mM IPTG for 3 hours at 37°C. Bacteria were then pelleted and resuspended in 20 mM sodium acetate pH 5.1 (pH adjusted with acetic acid). Resuspended cells were lysed by sonication and the lysate was clarified by centrifugation at 13,000 rcf, 4°C, 30 minutes. Clarified lysate was loaded onto a HiPrep^TM^ SP XL 16/10 cation exchange column (GE) preequilibrated in 20 mM sodium acetate pH 5.1. Column was washed with 5 column volumes of 20 mM sodium acetate pH 5.1 and then eluted with a gradient of 20 mM sodium acetate to 500 mM sodium acetate pH 5.1. The peak corresponding to WT ubiquitin was collected and further purified by size exclusion on an S75 16/60 column (GE) preequilibrated with 50 mM Tris pH 7.5, 150 mM NaCl. Peak corresponding to WT ubiquitin was collected, and quantified by UV-Vis absorption at 280 nm before flash freezing to store at −80°C for future use.

#### Preparation of Barnase

*E. coli* BL21 Rosetta 2 (DE3) cells were transformed with pEC099. Cells were then grown in 2 L LB Broth (Fisher) to 0.4 < OD_600_ < 0.8 and induced with 1 mM IPTG for 3 hours at 37°C. Bacteria were then pelleted and resuspended in 50 mM HEPES pH 7.0, 150 mM NaCl, 0.5 mM TCEP supplemented with 1X Halt protease inhibitor cocktail (Thermo) and benzonase (Novagen). Resuspended cells were lysed by sonication and the lysate was clarified by centrifugation at 13,000 rcf, 4°C, 30 minutes. The substrate was first purified by Ni^2+^-NTA affinity chromatography using its N-terminal His_6_. Clarified lysate was allowed to batch bind to HisPur^TM^ Ni^2+^-NTA resin (Thermo) washed with 50 mM HEPES pH 7.0, 150 mM NaCl, 25 mM imidazole, 0.5 mM TCEP and eluted with 50 mM HEPES pH 7.0, 150 mM NaCl, 500 mM imidazole, 0.5 mM TCEP. HRV3C-protease was added and the cleavage reaction was allowed to proceed overnight at 4°C under dialysis to 50 mM HEPES pH 7.0, 150 mM NaCl, 0.5 mM TCEP. HRV3C-protease and His-MBP tags were removed using a subtractive Ni^2+^-NTA purification step. Flow through was further purified by size exclusion chromatography using a S75 16/60 column (GE). Peak corresponding to barnase was collected, and quantified by UV-Vis absorption at 280 nm before addition of 10% glycerol and flash freezing to store at −80°C for future use.

#### Preparation of Ubiquitination Enzymes

Ubiquitination machinery *M. musculus* mE1, *S. cerevisiae* Ubc4, and *S. cerevisiae* Rsp5 were purified as described previously using the same procedure^15, 16^. *E. coli* BL21 Rosetta 2 (DE3) pLysS cells were transformed with pAM235 (mE1) or pAM236 (Ubc4) or pAM237 (Rsp5) and grown at 37°C in 6L of terrific broth (Novagen) until OD_600_ = 0.8 before expression was induced with 1 mM IPTG and allowed to continue overnight at 18°C. Cells were resuspended in 50 mM HEPES pH 7.6, 250 mM NaCl supplemented with protease inhibitors (pepstatin A, aprotonin, PMSF, and leupeptin), benzonase, and lysozyme (2 mg/mL) and stored at −80°C. Resuspended cells were thawed and lysed by sonication before lysate was clarified by centrifugation at 15,000 rcf for 30 mins at 4°C. Clarified lysate was batch bound to HisPur^TM^ Ni^2+^-NTA resin (ThermoFisher) equilibrated with 50 mM HEPES pH 7.6, 250 mM NaCl for one hour at 4°C. Resin was washed in a gravity flow column with at least 50 mL of 50 mM HEPES pH 7.6, 250 mM NaCl, 20 mM imidazole before protein was eluted with 50 mM HEPES pH 7.6, 250 mM NaCl, 250 mM imidazole. Eluate was concentrated in an Amicon spin concentrator (Millipore) and loaded onto a Superdex200 16/60 size exclusion column (GE) equilibrated in 20 mM HEPES pH 7.6, 100 mM NaCl, 10% glycerol. Peak corresponding with target protein was collected, concentrated in Amicon spin concentrator (Millipore), quantifying by absorbance at 28 0nm, and flash frozen in liquid nitrogen for storage at −80°C.

#### Preparation of AMSH Deubiquitinase

*E. coli* BL21 Rosetta 2 (DE3) pLysS cells were transformed with pAM241 and grown in 2 L of terrific broth (Novagen) at 37°C until OD_600_ = 0.6 after which expression was induced with 0.5 mM IPTG overnight at 18°C. Cells were resuspended in 50 mM HEPES pH 7.6, 250 mM NaCl supplemented with protease inhibitors (pepstatin A, aprotonin, PMSF, and leupeptin), benzonase, and lysozyme (2 mg/mL) and stored at −80°C. Resuspended cells were thawed and lysed by sonication before lysate was clarified by centrifugation at 15,000 rcf for 30 mins at 4°C. Clarified lysate was batch bound to HisPur^TM^ Ni^2+^-NTA resin (ThermoFisher) equilibrated with 50 mM HEPES pH 7.6, 250 mM NaCl for one hour at 4°C. Resin was washed with 50 mM HEPES pH 7.6, 250 mM NaCl, 10 mM ATP (to remove contaminating DnaK), 20 mM imidazole. The His_6_ tag was cleaved from AMSH by HRV3C-protease overnight at 4°C and AMSH was clarified through an ortho Ni^2+^-NTA step using HisPur Ni^2+^-NTA resin (ThermoFisher). Protein was concentrated in Amicon spin concentrator (Millipore) before being loaded on a S75 16/60 size exclusion column (GE) equilibrated with 20 mM HEPES pH 7.6, 100 mM NaCl, 10% glycerol. Peak corresponding to AMSH was collected, concentrated in an Amicon spin concentrator (Millipore), quantified by absorbance at 280 nm, and flash frozen in liquid nitrogen for storage at −80°C.

#### Preparation of homogenous mono-ubiquitinated substrate proteins

Substrate proteins, ubiquitin, ubiquitination enzymes, and AMSH were prepared as described above. Ubiquitination reactions were set up in reaction buffer (50 mM HEPES pH 8.0, 150 mM NaCl, 5 mM MgCl_2_, and 5% glycerol) in 20 μL aliquots as follows: 5 μM Uba1 (E1), 5 μM Ubc4 (E2), 5 μM Rsp5 (E3), 20 μM substrate, 750 μM WT ubiquitin or methylated ubiquitin, 5 mM ATP and incubated in a thermocycler for 3 hours at 25°C. 48 individual 20 μL reactions were performed for a typical prep. After three hours, reactions were pooled and HRV3C-protease was added and allowed to cleave overnight at 4°C. If WT ubiquitin was used, reactions were then treated with 0.5 μM AMSH for 30 minutes at room temperature and quenched with 5 mM EDTA. His-tagged ubiquitination machinery and the His-MBP scaffold were then removed via a subtractive Ni^2+^-NTA affinity step using a 1 mL HisTrap HP column (GE) pre-equilibrated with 50 mM HEPES pH 7.0, 150 mM NaCl, 25 mM imidazole. Flow through was then concentrated and loaded onto an S75i 10/300 size exclusion column (GE) pre-equilibrated with 25 mM HEPES pH 7.5, 150 mM KCl, and 15 mM MgOAc. The peak corresponding to the mono-ubiquitinated substrate was collected, concentrated, and quantified by UV-Vis absorption at 280 nm and 495 nm according to the manufacturer’s instructions before addition of 10% glycerol and flash freezing to store at −80°C for future use.

#### Preparation of mono-ubiquitinated substrate proteins with C-terminal ssrA tag/engageable tail

Substrate proteins and ubiquitination enzymes were prepared as described above. Ubiquitination reactions were set up in reaction buffer (50 mM HEPES pH 8.0, 150 mM NaCl, 5 mM MgCl_2_, and 5% glycerol) in 20 μL aliquots as follows: 5 μM Uba1 (E1), 5 μM Ubc4 (E2), 5 μM Rsp5 (E3), 20 μM substrate, 500 μM methylated ubiquitin (Millipore), 5 mM ATP and incubated in a thermocycler for 3 hours at 25°C. 24 individual 20 μL reactions were performed for a typical prep. After three hours, reactions were pooled and HRV3C-protease was added and allowed to cleave for 30 minutes at room temperature. Ubiquitination enzymes and His-MBP were removed by batch binding to MagneHis^TM^ (Promega) magnetic Ni^2+^-NTA resin for 1 hour at 4°C. Resin was pelleted in a magnetic tube rack, and the supernatant was collected for gel based single-turnover ubiquitin-independent degradation assays.

#### Preparation of Proteasome Lid Subcomplex

Lid subcomplex was recombinantly expressed and purified as described previously^16^. *E. coli* BL21-star(DE3) (Invitrogen) cells were transformed with pAM80, pAM85, and pAM86 for lid. pAM80 encodes for Sem1 and rare tRNA codons, pAM85 encodes Rpn5, MBP-HRV3C-Rpn6, Rpn8, Rpn11, and Rpn9, and pAM86 encodes Rpn3, His_6_-HRV3C-Rpn12, and Rpn7. Cells were grown in 2 L of terrific broth (Novagen) at 37°C until 1.0 < OD_600_ < 1.5 after which expression was induced with 1mM IPTG at 16°C for overnight. Bacteria were pelleted and resuspended in 60 mM HEPES pH 7.6, 50 mM NaCl, 50 mM KCl, 10 mM MgCl_2_, 5% glycerol and supplemented with protease inhibitors (aprotonin, pepstatinA, leupeptin, and PMSF or AEBSF), benzonase (Novagen), and 2 mg/mL lysozyme and stored at −80°C. Resuspended cells were lysed by sonication and the lysate was clarified by centrifugation at 21,000 rcf, 4°C, 30 minutes. Lid was first purified by Ni^2+^-NTA affinity chromatography via His_6_-HRV3C-Rpn12 using a 5mL HisTrap HP (GE) column, washed with 60 mM HEPES pH 7.6, 50 mM NaCl, 50 mM KCl, 10 mM MgCl_2_, 5% glycerol, 20 mM imidazole and eluted with 60 mM HEPES pH 7.6, 50 mM NaCl, 50 mM KCl, 10 mM MgCl_2_, 5% glycerol, 250 mM imidazole. Eluate was further purified via MBP-HRV3C-Rpn6 and amylose resin (NEB) and eluted with 60 mM HEPES pH 7.6, 50 mM NaCl, 50 mM KCl, 10 mM MgCl_2_, 5% glycerol, 10 mM maltose. Amylose eluates were cleaved with HRV3C-protease overnight at 4C before being loaded onto a Sup6i 10/300 size exclusion column (GE) pre-equilibrated with 60 mM HEPES pH 7.6, 50 mM NaCl, 50 mM KCl, 10 mM MgCl_2_, 5% glycerol, 0.5 mM TCEP. Peak corresponding to fully assembled lid was collected, concentrated, and quantified by UV-Vis spectroscopy before being flash frozen and stored at −80°C for future use.

#### Preparation of Proteasome Base Subcomplex and SspB_2_-fused Base Subcomplex

Base subcomplex was recombinantly expressed and purified as described previously^61^. *E. coli* BL21-star(DE3) (Invitrogen) cells were transformed with pAM81, pAM83, and pAM82 for wild-type base or pAM81, pAM83, and pAM210 for SspB_2_-Rpt2 base. pAM82 encodes for Rpt1, Rpt2, Rpt3, Rpt4, Rpt5, and Rpt6, pAM210 encodes Rpt1, SspB_2_-Rpt2, Rpt3, Rpt4, Rpt5, and Rpt6, pAM81 encodes Rpn1, Rpn2, and Rpn13, and pAM83 encodes rare tRNA codons and base chaperones (Nas6, Nas2, Rpn14, and Hsm3). Cells were grown in 3 L of terrific broth (Novagen) at 37°C until 0.6 < OD_600_ < 0.8 after which expression was induced with 1mM IPTG at 30°C for 5 hours followed by 16°C overnight expression. Bacteria were pelleted and resuspended in 60 mM HEPES pH 7.6, 50 mM NaCl, 50 mM KCl, 10 mM MgCl_2_, 5% glycerol, 1 mM ATP and supplemented with protease inhibitors (aprotonin, pepstatinA, leupeptin, and PMSF or AEBSF), benzonase (Novagen), and 2 mg/mL lysozyme and stored at −80°C. Resuspended cells were lysed by sonication and the lysate was clarified by centrifugation at 15,000 rcf, 4°C, 30 minutes. Base was first purified by Ni^2+^-NTA affinity chromatography via His_6_-Rpt6 using a 5mL HisTrap HP (GE) column, washed with 60 mM HEPES pH 7.6, 50 mM NaCl, 50 mM KCl, 10 mM MgCl_2_, 5% glycerol, 1 mM ATP, 20 mM imidazole and eluted with 60 mM HEPES pH 7.6, 50 mM NaCl, 50 mM KCl, 10 mM MgCl_2_, 5% glycerol, 1 mM ATP, 250 mM imidazole. Eluate was further purified via FLAG-Rpt1 and anti-FLAG M2 affinity resin (Sigma) and eluted with 60 mM HEPES pH 7.6, 50 mM NaCl, 50 mM KCl, 10 mM MgCl_2_, 5% glycerol, 1 mM ATP, 0.15 mg/mL FLAG peptide (Genscript). FLAG eluates were loaded onto a Sup6i 10/300 size exclusion column (GE) pre-equilibrated with 60 mM HEPES pH 7.6, 50 mM NaCl, 50 mM KCl, 10 mM MgCl_2_, 5% glycerol, 1 mM ATP, 0.5 mM TCEP. Peak corresponding to fully assembled base was collected, concentrated, and quantified by Bradford assay (BioRad) (source) using BSA (Sigma) as a standard before being flash frozen and stored at −80°C for future use.

#### Preparation of Proteasome Core Particle

20S core particle from *S. cerevisiae* was purified as described previously^62^ from yeast strain yAM54 bearing 3X-FLAG-Pre1. yAM54 cells were grown in 3 L of YPD at 30°C until saturation (3 days). Cells were pelleted and resuspended in 60 mM HEPES pH 7.6, 500 mM NaCl, 10 mM MgCl_2_, 5% glycerol, plunged into liquid nitrogen and subsequently stored at −80°C. Frozen resuspended cells were lysed using a 6875 Freezer Mill Dual Chamber Cryogenic grinder (SPEX Sample Prep). Lysate was diluted in 60 mM HEPES pH 7.6, 500 mM NaCl, 10 mM MgCl_2_, 5% glycerol and clarified by centrifugation at 15,000 rcf, 4C, 45 minutes. Base was first purified by anti-FLAG affinity chromatography using anti-FLAG M2 affinity resin (Sigma), exhaustively washed with 60 mM HEPES pH 7.6, 500 mM NaCl, 10 mM MgCl_2_, 5% glycerol, and eluted with 60 mM HEPES pH 7.6, 500 mM NaCl, 10 mM MgCl_2_, 5% glycerol, 0.15 mg/mL FLAG peptide (Genscript). Eluate was cleaved loaded onto a Sup6i 10/300 size exclusion column (GE) pre-equilibrated with 60 mM HEPES pH 7.6, 50 mM NaCl, 50 mM KCl, 10 mM MgCl_2_, 5% glycerol, 0.5 mM TCEP. Peak corresponding to fully assembled core was collected, concentrated, and quantified by UV-Vis spectroscopy before being flash frozen and stored at −80°C for future use.

#### Determination of global substrate stability by intrinsic tryptophan fluorescence

Two 5 μM protein stocks were prepared: A no denaturant protein stock and a high urea protein stock both in 25 mM HEPES pH 7.5, 150 mM KCl, and 15 mM MgOAc. The exact urea concentration in the high denaturant stock was determined by taking the refractive index. Samples with a range of urea concentrations were prepared by serial dilution of the two stocks and allowed to equilibrate at room temperature overnight. Measurements were then performed at 25°C using a PTI Quantamaster Fluorometer (Horiba). Tryptophan fluorescence was excited at 295 nm and a 10 second kinetic read of fluorescence emission at both 330 nm and 350 nm was performed at each denaturant concentration. Samples were recovered from the cuvette after each measurement and the exact urea concentration was determined by taking the refractive index. The signal was averaged over each 10 second period and reported as a ratio of average signal 330/average signal 350. Ratios were then normalized using equation 1 and fit to a two state folding model (equation 2) using Igor Pro 7, which allowed determination of the C_m_, ΔG_unfolding_, and *m*-value.

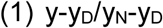

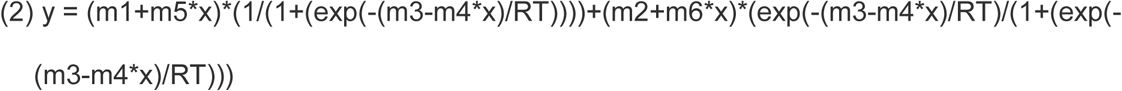

##### Parameter definitions

m1=folded intercept, m2 = unfolded intercept, m3 = ΔG_unfolding_, m4 = *m*-value, m5= folded baseline slope, m6=unfolded baseline slope

#### Determination of substrate native state energetics by native-state proteolysis

Ubiquitinated substrate sample prep was performed as described above except that AMSH deubiquitinase was allowed sufficient time to leave a mixed population of unmodified and mono-ubiquitinated species. Additionally, the final size exclusion step was omitted. Protein stocks were prepared in a 2 mL volumetric flask with final buffer of 25 mM HEPES pH 7.5, 150 mM KCl, and 15 mM MgOAc. Samples were allowed to equilibrate at room temperature in the dark overnight. Native State Proteolysis experimental protocol was adapted from previous work^36^. The equilibrated stock was divided into 200 μL aliquots and thermolysin protease (stock concentration 10 mg/mL) was added to a final concentration of 0.04 to 0.4 mg/mL. Time points (15 μL) were taken at (no protease control, 0:15, 0:30, 0:45, 1:00, 1:30, 2:00, 3:00, 5:00, 7:00, and 10:00) from the reactions and quenched in 2.5 μL of 0.5 M EDTA. 2.5 μL of 6X SDS-PAGE loading buffer was added to each sample and time points were run out on a 12% NuPAGE Bis-Tris^TM^ gel (Invitrogen) in 1X MES running buffer (50 mM MES, 50 mM Tris Base, 0.1% SDS, 1 mM EDTA). Gels were imaged using a BioRad ChemiDoc^TM^ and color inverted using the “Invert” command in ImageJ for ease of viewing and analysis. Band intensities of the unmodified and mono-ubiquitinated substrate bands were then quantified using ImageJ. SH3 and mono-ubiquitinated SH3 gels were quantified in ImageQuant (GE Healthcare) with a rolling ball background subtraction because proteolysis products could comigrate near full length protein. Band intensities were normalized to the no protease lane and fit to a first order exponential (equation 3) using IgorPro 7 to calculate the observed proteolysis kinetics (*k*_obs_). For a given substrate, *k*_obs_ was determined at several thermolysin concentrations and plotted against protease concentrations. ΔΔG_proteolysis_ was calculated from the slope of the linear fit to thermolysin vs. *k*_obs_. using equation 4 and equation 5. Individual ΔG_proteolysis_ could also be calculated using equation 6 and the measured *k*_cat_/K_M_ of thermolysin for a generic protein of 99,000 M^-1^s^-1^ ^36^.

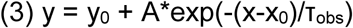

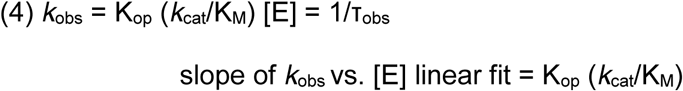

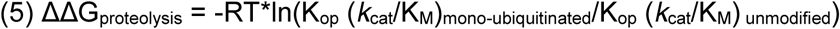

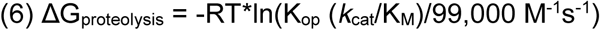

#### Cy5-methotrexate Binding to DHFR by Fluorescence Polarization

Equilibrium binding of Cy5-methotrexate to *M. smegmatis* DHFR was assessed by monitoring the increase in fluorescence polarization of Cy5-methotrexate upon binding DHFR. 50 nM Cy5-methotrexate was incubated with increasing concentration of unmodified or mono-ubiquitinated DHFR (quantified by fluorescein fluorescence using a standard curve) in 60 mM HEPES pH 7.6, 50 mM NaCl, 50 mM KCl, 10 mM MgCl_2_, 5% glycerol, 0.5 mM TCEP for 20 minutes at room temperature to reach equilibrium. Fluorescence polarization was monitored for 5 minutes on a Synergy Neo2 multi-mode plate reader. Time points were averaged and normalized to Cy5-methotrexate in the absence of DHFR. For the unmodified DHFR, K_d_ was determined by fitting the change in fluorescence polarization as a function of DHFR concentration to simple single site binding model^63^ (Equation 7).

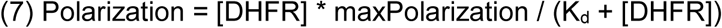

### Proteasome Degradation Assays

#### Preparation of Polyubiquitinated Barstar Variants

Barstar ubiquitination was performed exactly as above except that AMSH removal of K63-linked polyubiquitin chains was omitted. Ortho-Ni^2+^ purified ubiquitinations were subsequently separated by size-exclusion chromatography on an S200i 10/300 (GE Healthcare) and 0.5 mL fractions were assessed for degradable species by incubating with proteasome under single turnover conditions at 30°C for 30 minutes and analyzing products by SDS-PAGE (Supplementary Fig. 4b).

#### Gel Based Single-Turnover Ubiquitin-Independent Degradation Assay

2X stocks of substrate (300 nM final) were prepared in assay buffer (60 mM HEPES pH 7.6, 50 mM NaCl, 50 mM KCl, 10 mM MgCl2, 0.5 mM TCEP, 5 mM ATP, 5% glycerol, 1 mg/ml BSA). 2X proteasome stocks were performed by reconstituting recombinant lid (5 μM final), recombinant SspB_2_-Rpt2 base (5 μM final), recombinant Rpn10 (5 μM final), and core particle (2.5 μM final) in assay buffer with an ATP-regeneration system (creatine kinase, creatine phosphate, and 5 mM ATP) and allowed to assemble for 3 minutes at room temperature. Reactions were performed in technical triplicate at 30°C in a thermocycler and initiated by mixing equivolume (12.5 μL) of 2X substrate with 2X proteasome. Time points (1.2 μL) were taken at (0:10, 0:20, 0:30, 0:45, 1:00, 1:30, 2:00, 3:00, 5:00, 10:00, 15:00, 20:00, 30:00 min) from the reactions and quenched in 5 μL of 2X SDS-PAGE loading buffer (125 mM TrisHCl pH 6.8, 20% glycerol, 4% SDS). Gel samples were separated by electrophoresis on 4-20% TGX gels (Bio-Rad) before fluorescence imaging on a typhoon variable mode scanner (GE) with 50 μm per pixel density. Images were quantified in ImageQuant (GE) by normalizing band intensity of each species per total lane intensity to account for loading variation. Quantified species were plotted as percent total signal (Supplementary Fig. 3c) and fit to a single exponential equation (Equation 3) in IgorPro7. For degradations performed with ATPγS, proteasomes were assembled in ATP for 3 minutes at room temperature, then ATPγS was added (5 mM final) for 3 minutes at room temperature prior to substrate addition. For degradations using only the core particle, core particle was added to 900 nM final with substrate and incubated at 30°C for the indicated time points. Time points of 0, 10:00, and 30:00 minutes were quenched in SDS-PAGE loading buffer for trials involving core particle only or ATPγS inhibited proteasome and separated by SDS-PAGE on 4-20% and assess qualitatively.

#### Fluorescence Polarization Single-Turnover Ubiquitin-Independent Degradation Assay

For ubiquitin-independent degradations assessed by fluorescence polarization, reactions were initiated with equivolume (2.5 μL) addition of substrate to proteasome directly within a 384-well black bottom plate (Corning) and fluorescence polarization was monitored in a Synergy Neo2 multimode plate reader (BioTek). Decreased fluorescence polarization over time as substrate was processed into peptides could also be fit to a single exponential model (Equation 3) in IgorPro7.

#### Fluorescence Polarization Single-Turnover Ubiquitin-Dependent Degradation Assay of Ub_n_ Barstar Variants

Substrates were prepared to 2X concentration (6 nM final) in assay buffer. Proteasome was reconstituted to 2X concentration in assay buffer (2.5 μM lid, base, and Rpn10 with 0.9 μM core particle) and allowed to assemble for 3 minutes at room temperature prior to reaction initiation. Reactions were initiated with appropriate dilution of 2X substate (2.5 μL) into 2X proteasome (2.5 μL) in a 384-well black bottom plate (Corning) and the decrease of fluorescence polarization over time was monitored on a Synergy Neo2 multimode plate reader (BioTek). Trials were repeated for n=3 and averages collated with standard deviations. Where exponential decay was observed, curves could be fit to a single exponential model (Equation 3) in IgorPro7. For reactions performed with core particle only, core particle was made to 2X concentration (1.8 μM) and added equivolume with 2X substrate (5 μL final) and fluorescence polarization was monitored as above. Single turnover conditions were verified by single reactions with doubled proteasome concentration by reconstituting proteasome to 4X concentration and diluting with equivolume substrate (2.5 μl each) to 2X proteasome and monitoring fluorescence polarization kinetics as described above. For degradations with *o*-phenanthroline inhibited proteasomes, proteasomes were allowed to assemble at 3X concentration for 3 minutes at room temperature between dilution with *o-*phenanthroline (30 mM stock in assay buffer; 5 mM final) to 2X concentration for 2 minutes before degradation initiation as described above. For degradations using only the core particle, core particle was added to 900 nM final with substrate as described above.

#### Fluorescence Polarization Single-Turnover Ubiquitin-Dependent Degradation Assay of Substrates in the Presence of Barnase

Substrates were prepared to 2X concentration (6 nM final) in assay buffer with barnase added in excess (20 μM final) and allowed to come to equilibrium for greater than 5 minutes at room temperature^63^ prior to degradation initiation. Degradations were performed exactly as described above. Saturation of barnase binding was assessed by doubling barnase concentration (40 μM final) and comparing fluorescence polarization kinetic differences. FAM-Titin-I27^V13P,V15P^ ubiquitinated as described above was degraded in the presence or absence of 20 μM barnase with proteasome at the same concentration as described.

---

**Supplementary Fig. 1.**
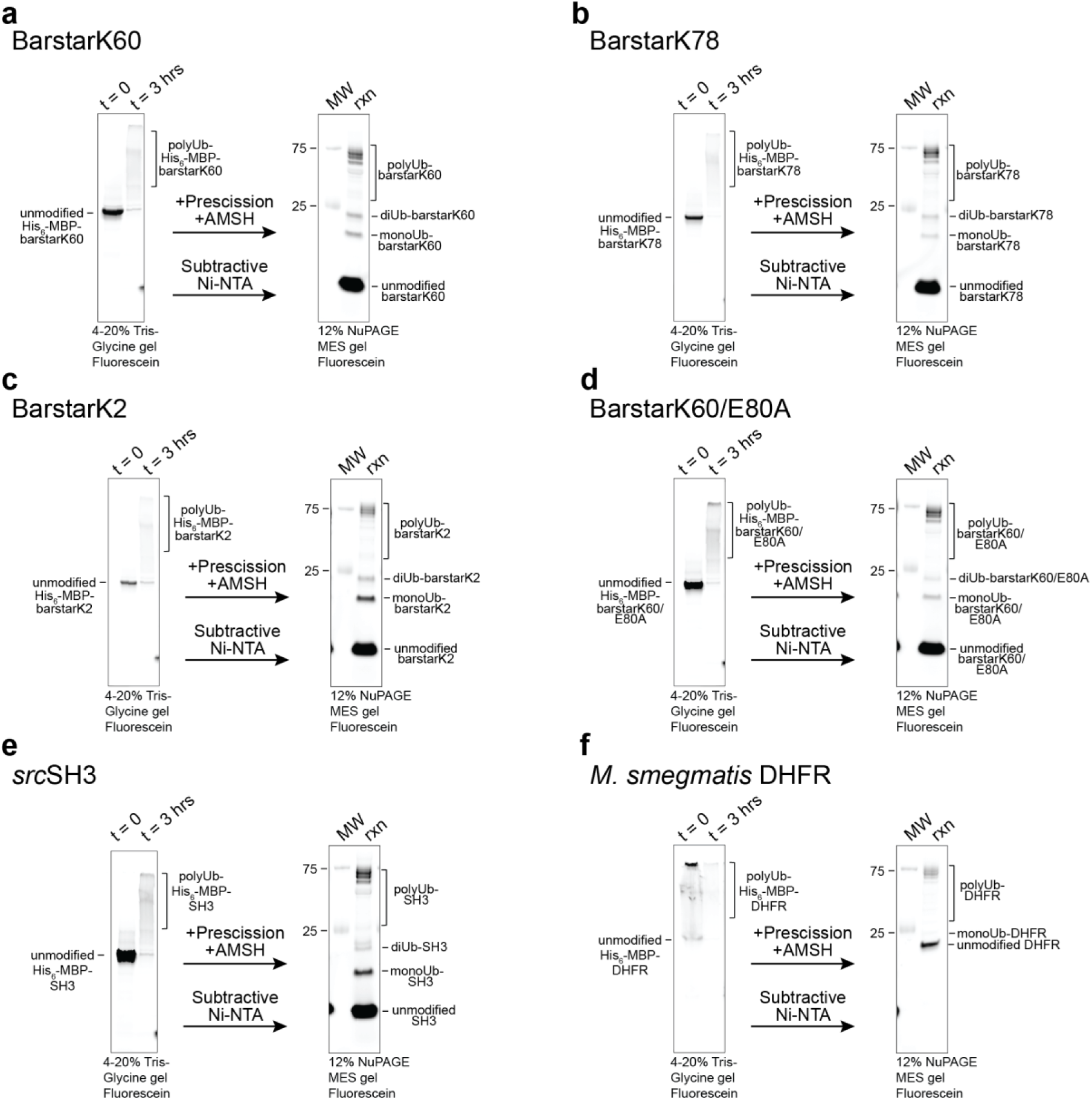
Biochemically reconstituted *in vitro* ubiquitination system for ubiquitination of diverse single-lysine substrates. **(a-f)** Fluorescence scans of SDS-PAGE gels showing the full-length substrates immediately after reaction initiation and again after 3 hours of ubiquitination. Reactions were treated with Prescission (HRV3C) protease and AMSH deubiquitinase prior to subtractive Ni^2+^-NTA chromatography to reveal clearly defined mono- and di-ubiquitinated substrate with native isopeptide linkages.

**Supplementary Fig. 2.**
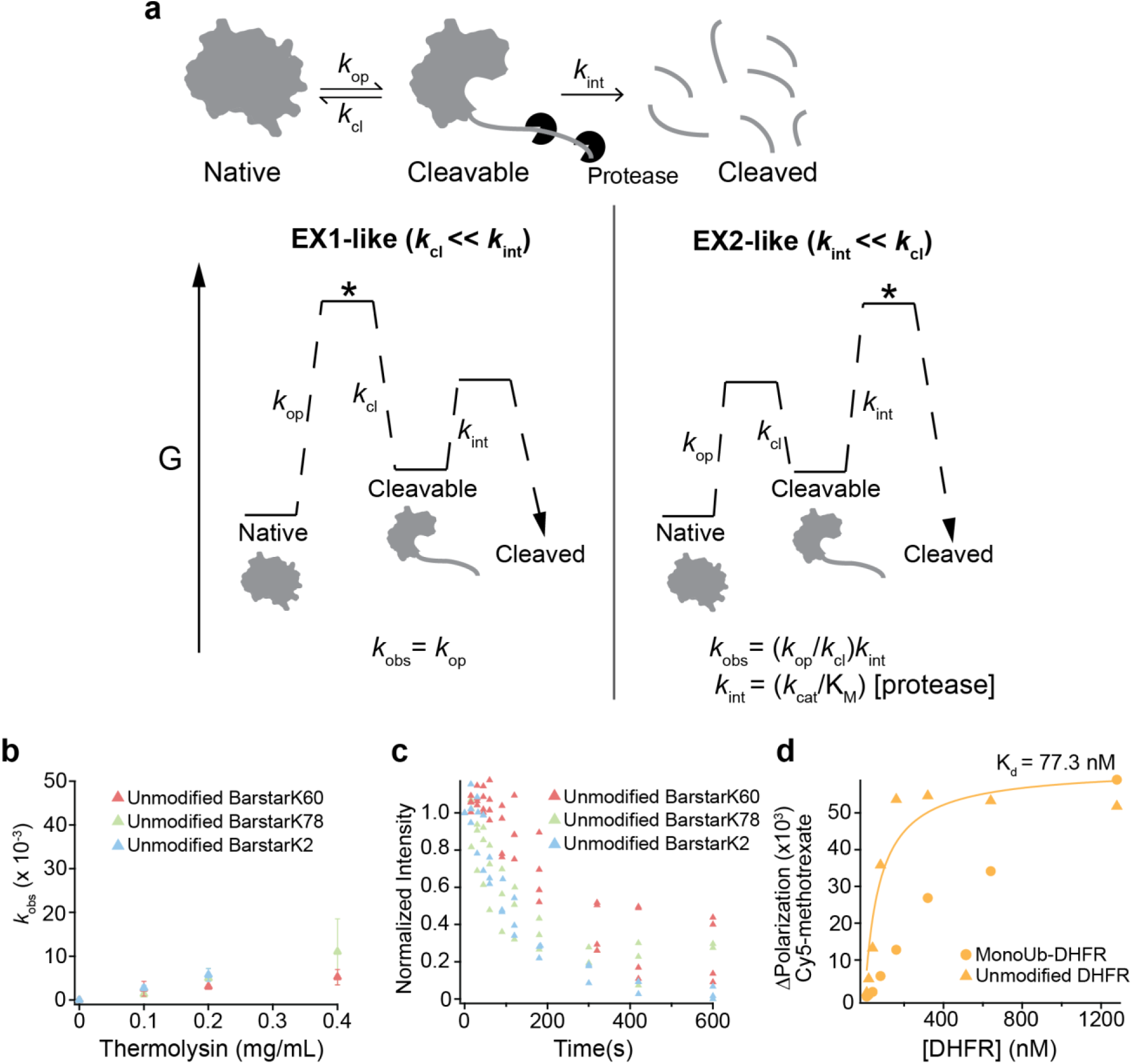
Explanation of native state proteolysis for investigating the energetics of partial unfolding and validation of native state proteolysis with these systems. **(a)** EX1 and EX2-like kinetic regimes for native state proteolysis. In EX1-like experiments, the conformational change between the native and cleavable states is the rate-limiting step, and *k*_obs_ = *k*_op_. In EX2-like experiments, as shown in Fig. 2, the proteolysis of the cleavable state is the rate-limiting step, and *k*_obs_ = K_op_(*k*_cat_/K_M_)[protease]. **(b)** Linear fit of thermolysin concentration vs. *k*_obs_ for unmodified barstarK2, K60, and K78 showing identical proteolysis kinetics at all concentrations of thermolysin used. **(c)** Example exponentials for unmodified barstarK2, K60, and K78 for multiple replicates at 0.2 mg/mL thermolysin showing the similarity of *k*_obs_ for the three unmodified barstar variants. **(d)** Binding curve for unmodified *M. smegmatis* DHFR and monoUb-*M. smegmatis* DHFR to Cy5-methotrexate measured by Cy5 fluorescence polarization.

**Supplementary Fig. 3.**
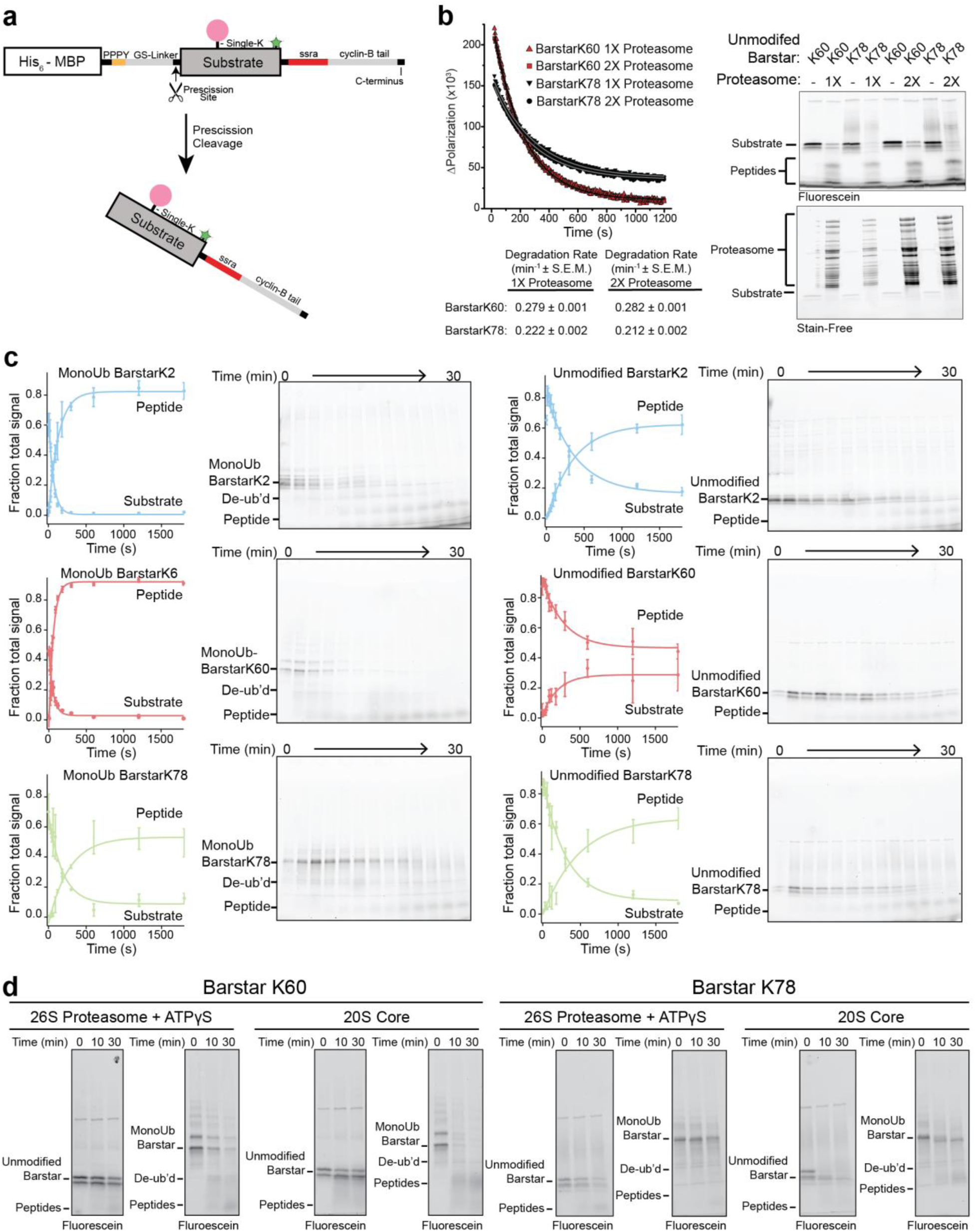
Ubiquitin-independent substrate delivery allows for comparison of unmodified with mono-ubiquitinated barstar variants. **(a)** Schematic showing substrate design for ubiquitin-independent delivery whereby *in vitro* ubiquitination system as in Fig. 1a is used for enzymatic ubiquitin ligation of methylated ubiquitin to a single lysine on barstar containing a C-terminal unstructured region with an ssrA tag, zero-lysine cyclin-B tail and C-terminal zero-lysine Strep(II) tag for selection. Scaffolding was removed by Prescission (HRV3C) protease and a subtractive Ni^2+^-NTA affinity step. **(b)** Confirmation of single turnover degradation conditions via proteasome doubling. BarstarK60 and barstarK78 degradation were monitored by fluorescence polarization over time with proteasome at the indicated concentrations. Observed rates are reported with S.E.M. Right, end-point SDS-PAGE gel of degradation showing conversion of barstarK60 and barstarK78 to peptides (fluorescein channel) and total protein estimated by Stain-Free imaging (Bio-Rad). **(c)** Representative SDS-PAGE gels of ubiquitin-independent degradations with identified bands indicated at left for each variant. Quantifications and fits of substrate bands are left of the representative gels. “Substrate” and “Peptide” bands as a fraction of total lane intensity are presented as averages with standard deviations (n=3). **(d)** Fluorescence scans of SDS-PAGE gels from time courses of unmodified or monoUb-barstarK60 and barstarK78 degradations with proteasome in the presence of ATPγS or core particle alone. Identified bands are indicated.

**Supplementary Fig. 4.**
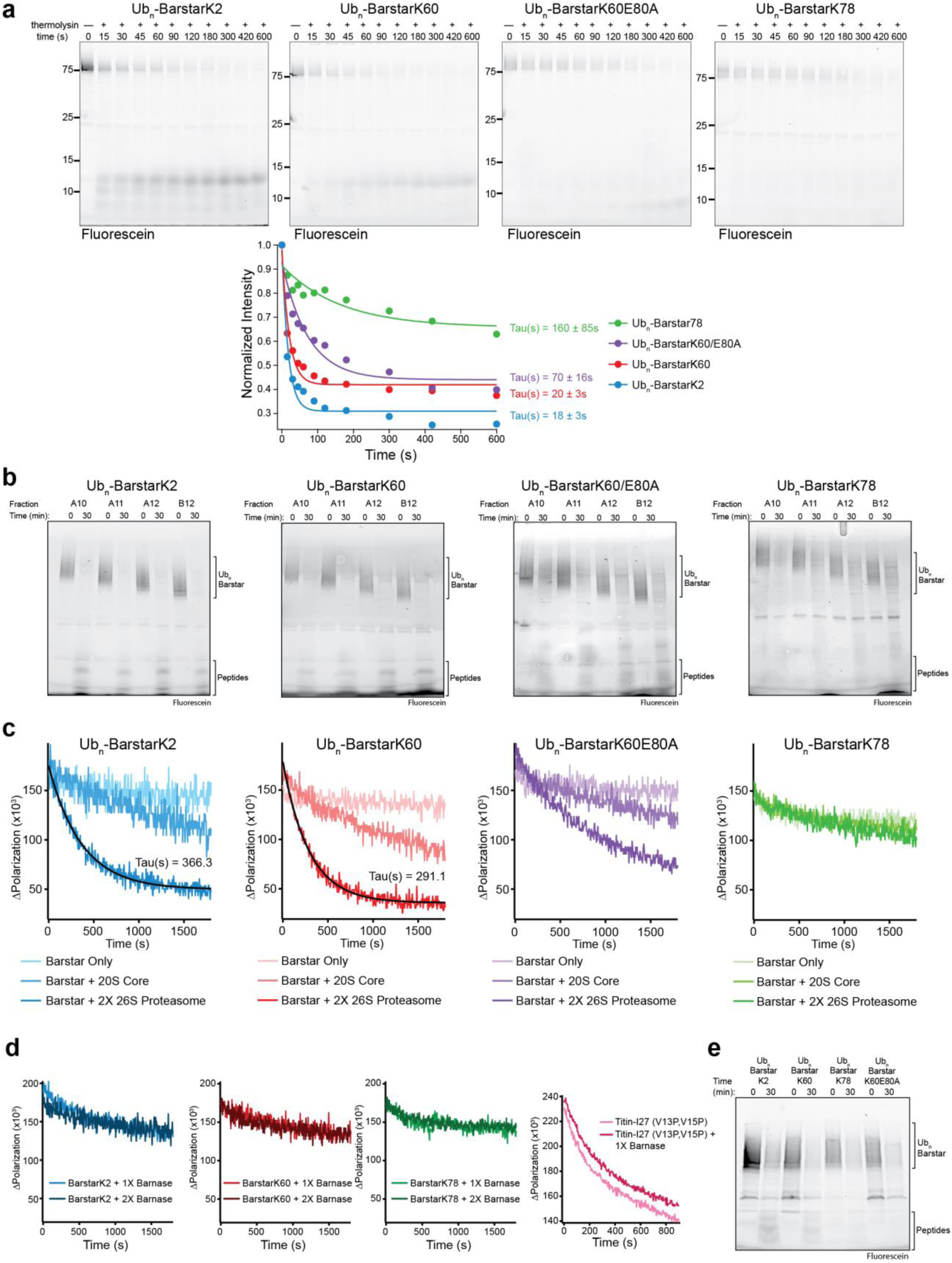
Ubiquitinated substrate energetics mediate proteasomal engagement and degradation. (**a**) Fluorescein scans of 12% Bis-Tris Nu-PAGE (Invitrogen) SDS-PAGE gels of thermolysin proteolysis overtime of Ub_n_-barstar variants with molecular weight standards and time points indicated (Top). Quantified gel bands were normalized and plotted against time and time constants are reported with S.E.M. (**b**) Fluorescein scan of 4-20% TGX (Bio-Rad) SDS-PAGE gels from end-point single turnover degradations of fractions obtained from size exclusion chromatography of Ub_n_-barstar variants with poly-ubiquitinated species and peptides indicated. Fraction A12 from each gel is presented in Fig. 4. (**c**) Fluorescence polarization of Ub_n_-barstar substrates treated with 2X concentration of proteasome, 900 nM of core particle, or untreated. Times constants are reported with S.E.M (**d**) Fluorescence polarization of single-turnover degradations of Ub_n_-barstars in the presence of 20 μM (1X) or 40 μM (2X) barnase (Left). Single-turnover degradation of FAM-Titin-I27^V13PV15P^ in the presence or absence of 20 μM (1X) barnase monitored by fluorescence polarization (Right). (**e**) Fluorescence scan of 4-20% TGX (Bio-Rad) SDS-PAGE gels from end-point reactions of Ub_n_-barstar variants with excess core particle (900 nM). Peptides and Ub_n_-barstar species are indicated.

**Supplementary Fig. 5.**
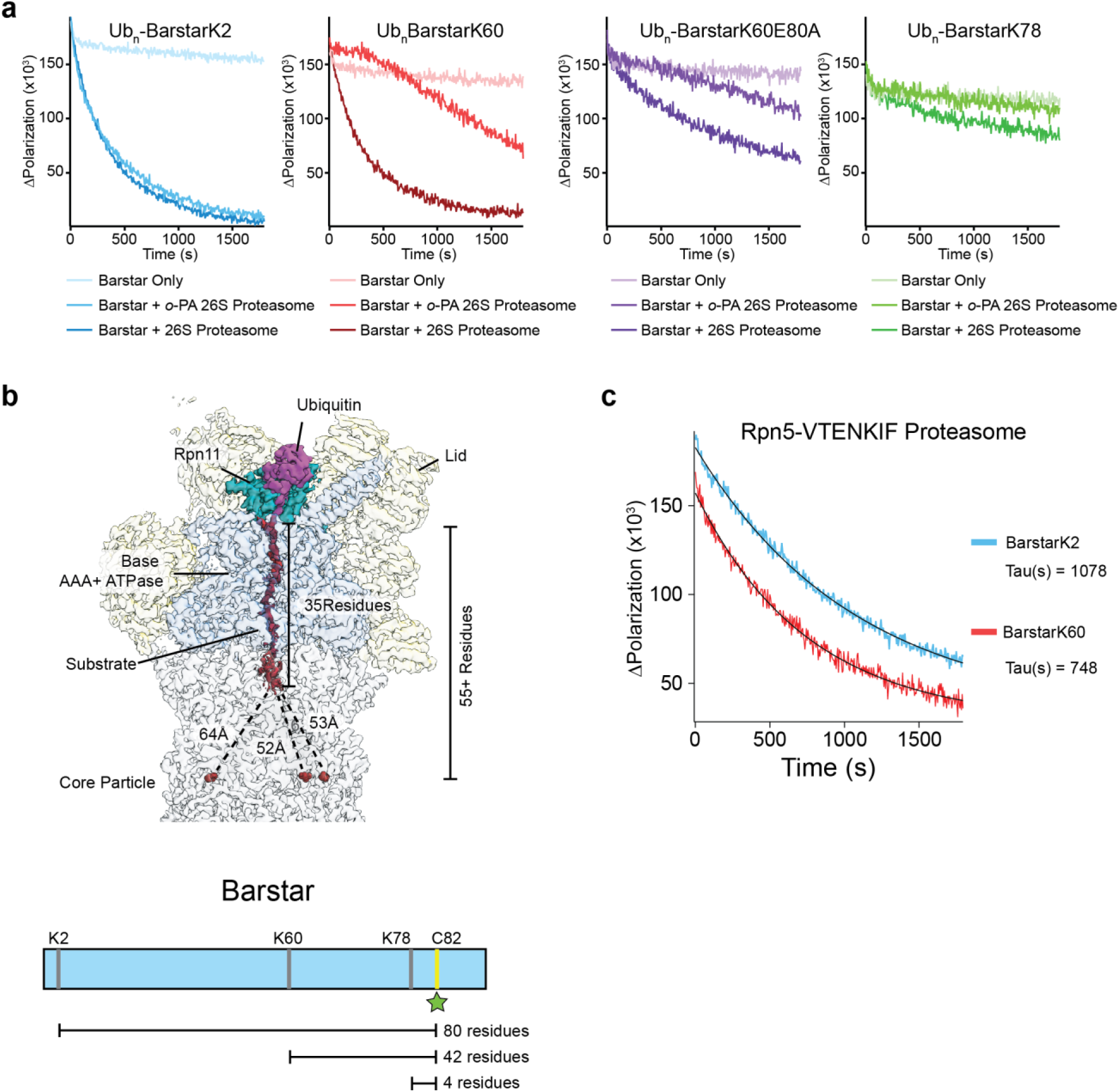
Proteasome engagement can be the rate limiting step for degradation. (**a**) Single-turnover degradations of Ub_n_-barstar substrates in the presence and absence of Rpn11 inhibitor *o*-phenanthroline monitored by fluorescence polarization. (**b**) Substrate bound proteasome density (EMD: 9045, PDB: 6FVW) with lid subunits and Rpn1 and Rpn2 in yellow, ubiquitin in magenta, Rpn11 in dark cyan, the base AAA+ ATPase in cornflower blue, substrate polypeptide in red, and the core particle in light grey. Distances were obtained from PDB: 6FVW. Below, cartoon of barstar sequence topology highlighting singe lysine positions and the single, fluorescein-labeled, cysteine at position 82. (**c**) Single-turnover degradations of Ub_n_-barstarK2 and Ub_n_-barstarK60 by Rpn5-VTENKIF proteasome and calculated time constants.

